# Novel Roles for Diacylglycerol in Synaptic Vesicle Priming and Release Revealed by Complete Reconstitution of Core Protein Machinery

**DOI:** 10.1101/2023.06.05.543781

**Authors:** R Venkat Kalyana Sundaram, Atrouli Chatterjee, Manindra Bera, Kirill Grushin, Aniruddha Panda, Feng Li, Jeff Coleman, Seong Lee, Sathish Ramakrishnan, Andreas M. Ernst, Kallol Gupta, James E. Rothman, Shyam S. Krishnakumar

**Author notes:** Correspondence (JER); (SSK). These authors contributed equally (R Venkat Kalyana Sundaram, Atrouli Chatterjee).

## Abstract

Here we introduce the full functional reconstitution of genetically-validated core protein machinery (SNAREs, Munc13, Munc18, Synaptotagmin, Complexin) for synaptic vesicle priming and release in a geometry that enables detailed characterization of the fate of docked vesicles both before and after release is triggered with Ca^2+^. Using this novel setup, we discover new roles for diacylglycerol (DAG) in regulating vesicle priming and Ca^2+-^triggered release involving the SNARE assembly chaperone Munc13. We find that low concentrations of DAG profoundly accelerate the rate of Ca^2+^-dependent release, and high concentrations reduce clamping and permit extensive spontaneous release. As expected, DAG also increases the number of ready-release vesicles. Dynamic single-molecule imaging of Complexin binding to ready-release vesicles directly establishes that DAG accelerates the rate of SNAREpin assembly mediated by Munc13 and Munc18 chaperones. The selective effects of physiologically validated mutations confirmed that the Munc18-Syntaxin-VAMP2 ‘template’ complex is a functional intermediate in the production of primed, ready-release vesicles, which requires the coordinated action of Munc13 and Munc18.

**SIGNIFICANCE STATEMENT:** Munc13 and Munc18 are SNARE-associated chaperones that act as “priming” factors, facilitating the formation of a pool of docked, release-ready vesicles and regulating Ca^2+^-evoked neurotransmitter release. Although important insights into Munc18/Munc13 function have been gained, how they assemble and operate together remains enigmatic. To address this, we developed a novel biochemically-defined fusion assay which enabled us to investigate the cooperative action of Munc13 and Munc18 in molecular terms. We find that Munc18 nucleates the SNARE complex, while Munc13 promotes and accelerates the SNARE assembly in a DAG-dependent manner. The concerted action of Munc13 and Munc18 stages the SNARE assembly process to ensure efficient ‘clamping’ and formation of stably docked vesicles, which can be triggered to fuse rapidly (∼10 msec) upon Ca^2+^ influx.

## INTRODUCTION

Information transfer in the brain depends on the controlled yet rapid (millisecond) release of neurotransmitters stored in synaptic vesicles (SVs). SV fusion is mediated by synaptic SNARE proteins, VAMP2 (v-SNARE) on the vesicles and Syntaxin/SNAP25 (t-SNAREs) on the target pre-synaptic membrane (1, 2). When an SV approaches the plasma membrane (PM), the helical SNARE motifs of the v- and t-SNARE progressively assemble (‘zipper’) into a ternary complex, which provides the energy to fuse the opposing membranes (3, 4). Neurotransmitter release is tightly controlled by pre-synaptic calcium (Ca^2+^) concentration and occurs from a readily-release pool (RRP) of vesicles that are ‘primed’ to rapidly fuse upon Ca^2+^-influx (5-8).

The ‘priming’ process involves a series of preparatory molecular reactions that properly aligns the v- and t-SNAREs for polarized assembly and arrest (‘clamps’) SNARE zippering in an intermediate state (9-12). This ensures that every RRP vesicle contains multiple partially-assembled ‘SNAREpins’, close to the point of triggering fusion, which can be synchronously released by Ca^2+^ to drive ultra-fast fusion (13). Vesicle priming is orchestrated by molecular chaperones (Munc13 and Munc18) which guide the SNARE assembly process, and regulatory elements (Synaptotagmin and Complexin) which act together to ‘clamp’ the SNARE assembly to produce RRP vesicles (14-17). The nucleation of the SNARE assembly has been shown to be error-prone and slow; thus, the combined action of specialized chaperones, Munc18 and Munc13 is crucial for efficient priming (16).

Munc18 forms a tight binary complex with Syntaxin, which is essential for proper Syntaxin folding and recruitment of the Munc18/Syntaxin complex to the active zone (18-20). This interaction also maintains Syntaxin in a ‘closed’ conformation, preventing premature binding with SNAP25 on the pre-synaptic membrane (15, 17, 21). Recent structural analyses, which revealed that Munc18 can simultaneously bind Syntaxin and VAMP2 SNARE motifs (22, 23), suggest that Munc18 may also serve as a template for nucleating SNARE assembly by holding Syntaxin and VAMP2 in proper alignment. The functional importance of the Munc18-Syntaxin-VAMP2 ‘template’ complex has been inferred by single-molecule force spectroscopy and bulk-fusion assays (24-26).

Munc13, a large multi-domain protein (21, 27-29), also plays a key and multi-faceted role in the SV priming process. Specifically, Munc13 catalyzes the SNARE nucleation by ‘opening’ the closed Syntaxin/Munc18 complex to enable the formation of the Munc18-Syntaxin-VAMP2 ‘template’ complex (16, 17, 21, 30). Munc13 also promotes its progression into a ‘ternary’ SNARE complex with SNAP25 (16). This catalytic function and associated SNARE interactions have been mapped to the large central module within Munc13 called the MUN domain (30). Additionally, Munc13 is believed to be involved in the initial tethering and docking of SV at the active zone by acting as a “molecular bridge” that links the SV and PM. This function is thought to involve interactions between the C1-C2B region and the C2C domain located on opposite ends of the MUN domain (17, 27, 31). The conserved C1-C2 domains have been shown to mediate interactions with lipids [e.g. phosphatidylserine (PS), phosphatidylinositol 4,5-bisphosphate (PIP2), diacylglycerol (DAG), Ca^2+^, and other proteins as a means of modulating Munc13 docking/priming function (17, 32-36). Accumulating evidence shows that activation of these domains, separately or in combination, has a profound stimulatory effect on the replenishment and priming of the RRP vesicles both during and after high neuronal activity (37, 38). Recently distinct oligomeric assemblies of Munc13 have been described which are likely to function at distinct stages of vesicle priming and to be differentially regulated by DAG (27, 36).

The precise roles of Munc13 and its ligand DAG in vesicle priming and release remain to be established. This is largely due to the overlapping nature of their functions and a lack of functional reconstitutions with sufficient resolution to dissect their contributions at different stages of vesicle docking, priming, and fusion. We have recently described a biochemically fully-defined experimental setup based on a suspended lipid membrane (SLIM) platform that is well-suited for this purpose (39-42). This high-throughput, cell-free platform can precisely track individual vesicle docking, priming (clamping) and Ca^2+^-triggered release events with millisecond resolution using fluorescence microscopy. Tracking of single vesicles allows for isolated measurements of vesicle docking, clamping/priming, and Ca^2+^-triggered fusion, independent of any alteration in the preceding or subsequent steps. When combined with Total Internal Reflection Fluorescence (TIRF) microscopy, it also enables the visualization of the dynamics and organization of individual proteins beneath the docked vesicles (41). The approach is highly versatile in that all critical components, including the identity and density of protein, and the composition of buffers and membranes, can be rigorously controlled and thus, allows us to build experimentally constrained molecular models which are tightly correlated with physiology through mutational effects. Indeed, we recently employed this *in vitro* setup to dissect the dual clamp/activator function of Synaptotagmin and Complexin in molecular terms and develop detailed mechanistic models (13, 42).

Here, using the new iteration of the *in vitro* single-vesicle fusion assay, which accurately reproduces the physiological start point for SNARE recruitment and priming, we report that DAG is an essential co-factor for Munc13 chaperone function and balanced activities of Munc18 and Munc13 are required for efficient SNARE priming and Ca^2+^-evoked vesicular release.

## RESULTS

### Reconstitution of Munc13-regulated vesicle docking and fusion

To dissect the molecular mechanisms underlying Munc13-1 function, we adapted our *in vitro* single-vesicle fusion assay (13, 40, 42) to directly investigate the Munc13-dependent vesicle docking and fusion process (Figure 1). To best approximate the physiological conditions, we reconstituted Syntaxin with its chaperone Munc18 as a 1:1 functional complex along with palmitoylated SNAP25 at a very low concentration (1:3200 protein: lipid ratio) in the suspended lipid membrane containing 15% PS and 3% PIP2 (SI Appendix, Figure S1) (43). We employed small unilamellar vesicles (SUVs) with an average of 74 copies (outward facing) of VAMP2 and 25 copies of Synaptotagmin-1 (outward facing) and included Complexin-1 (CPX, 2 µM) in the solution (SI Appendix, Figure S1). In all experiments, we used a conserved C-terminal portion of Munc13-1 consisting of the contiguous C1-C2B-MUN-C2C domains (Munc13-1 residues 529-1735; henceforth referred to as Munc13), except for residues 1408-1452 within a non-conserved loop in the MUN domain, which were deleted and replaced with EF residues to minimize dimerization and oligomerization (21, 44). We monitored large ensembles of vesicles (∼200) and used fluorescently labelled lipid (2% ATTO647N-PE), or a content dye (Sulforhodamine B) introduced in the SUVs to track the docking, un-docking, clamping and fusion (either spontaneous or Ca^2+^-evoked) of individual vesicles (SI Appendix Figure S2).

**Figure 1.**
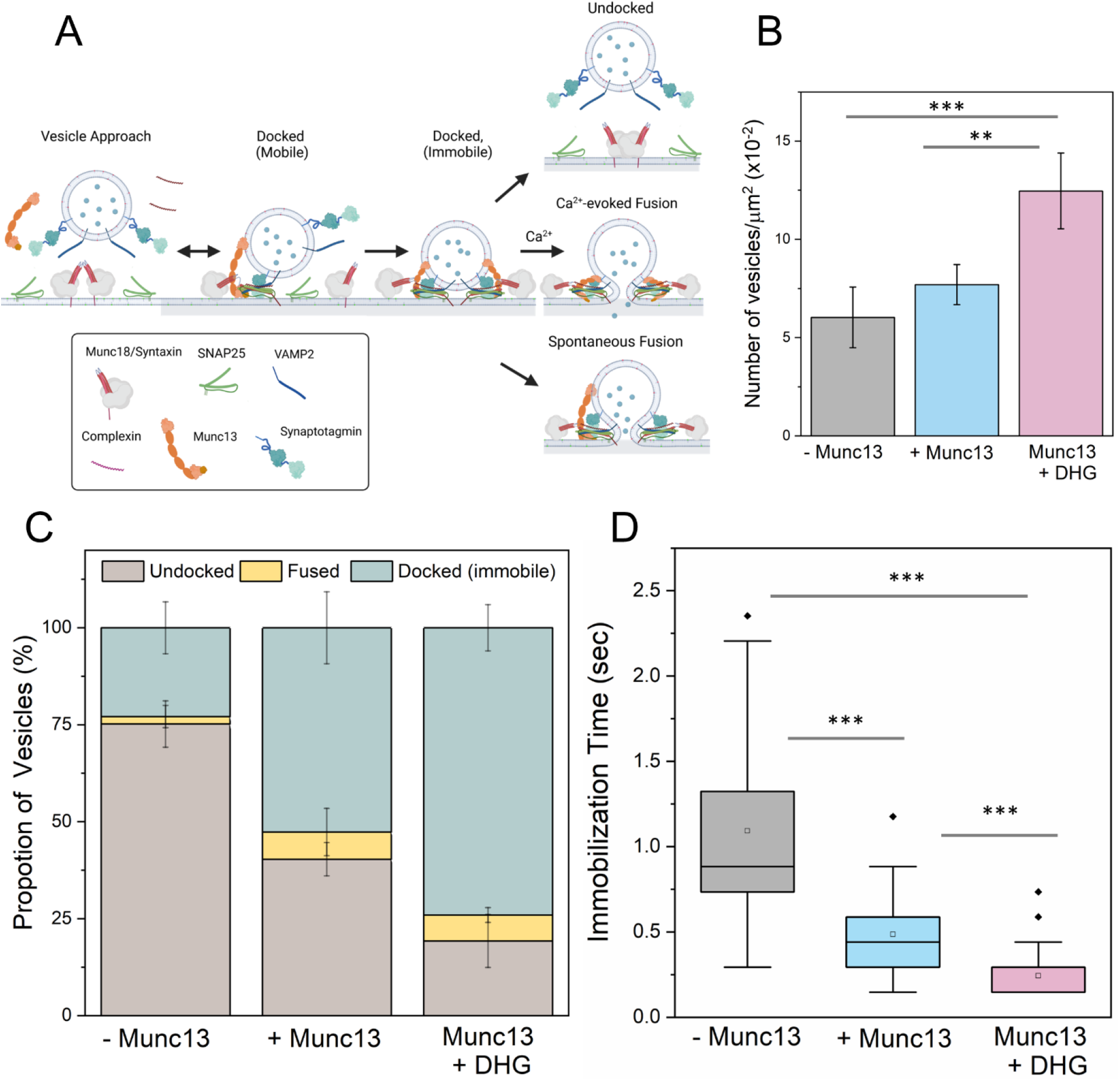
Munc13 and DHG promote the formation of stably, docked ‘immobile’ vesicles. A) To reconstitute the Munc13/Munc18-dependent vesicle priming and fusion process under cell-free conditions, we included a 1:1 Syntaxin/Munc18 complex and palmitoylated SNAP25 in the suspended lipid membrane, as well as VAMP2/Synaptotagmin in SUVs, with CPX and Munc13 in solution. We monitored the fate of a large number of individual vesicles (∼200 per condition) using fluorescent labels in the SUVs. (B) The number of vesicles attached or ‘docked’ to the suspended bilayer did not appreciably change without or with Munc13 (grey and blue bar respectively), but DHG activation (pink bar) of Munc13 significantly increased the number of docked vesicles. (C) The fate of the docked vesicles strongly depended on the availability of Munc13 and its activation by DHG. In the absence of Munc13 or DHG, most vesicles undock (brown bar), with only a small proportion converting into the ‘immobile’ docked stage (green bar). The inclusion of Munc13 increased the number of immobile vesicles, which was further enriched by the addition of DHG. In all cases, we observed a small percentage of spontaneous fusion events (yellow bar). (D) DHG and Munc13 also accelerated the formation of the immobile docked stage. Without Munc13 or DHG, the docked vesicles reached the immobile state over 1-2 seconds. The addition of Munc13 lowered this transition time to ∼0.5 seconds, which was further reduced by DHG to ∼0.25 seconds. The average values and standard deviations from 3-4 independent experiments (with ∼200 vesicles per condition) are shown. **p<0.05; *** p<0.005 using the student’s t-test.

In the absence of Munc13, most of the docked vesicles remained mobile on the bilayer surface, with the majority of vesicles undocking (∼79%) or fusing spontaneously (∼2%). Consequently, only a small fraction (∼19%) of vesicles converted into an ‘immobile’ (clamped) state and remained unfused during the initial 3 min observation window (Figure 1A-C). Adding Munc13 (200 nM) to the solution resulted in a small (∼20%) increase in the total number of docked vesicles (Figure 1B), but it significantly reduced the undocking of vesicles (∼40%). The level of spontaneous fusion during this period was low (∼8%) resulting in a large fraction (∼52%) of the vesicles remaining stably docked, immobile and unfused (Figure 1B, C). The addition of Munc13 also lowered the time the docked vesicles took to reach the immobile ‘clamped’ state from ∼1.1 sec to ∼500 msec (Figure 1D).

The addition of Ca^2+^ (100 µM) triggered the fusion of the stably, clamped vesicles (Figure 2) but to varying degrees depending on prior conditions. In the absence of Munc13, Ca^2+^ influx triggered fusion (as measured by lipid mixing at 147 msec frame rate) of ∼25% of docked immobile vesicles, while the inclusion of Munc13 increased the proportion of Ca^2+^-evoked fusion to ∼60% of the docked vesicles (Figure 2A). Munc13 also accelerated the fusion kinetics, with the majority fusing within 6 sec after Ca^2+^-addition with an estimated half-life (t1/2) of ∼2.2 secs (Figure 2B). In contrast, without Munc13, the vesicles fused slowly and intermittently over 10 secs (Figure 2B). In these experiments, we used a lipid-conjugated Ca^2+^ indicator (Calcium green C24) attached to the planar bilayer to estimate the time of arrival of Ca^2+^ at/near each docked vesicle (42). Overall, our data showed that the inclusion of Munc13 significantly improved the formation of stably docked vesicles, along with the probability and kinetics of Ca^2+^-evoked fusion. However, the extent of priming and efficiency of subsequent release for this Munc13-dependent reaction was much lower than when the requirement for Munc13 and Munc18 chaperones was by-passed by employing pre-assembled t-SNAREs, where >90% of vesicles remained stably docked and fused synchronously with a fusion rate <100 msec (13, 39, 42). This suggested that additional components might be crucial for optimal Munc13 function.

**Figure 2.**
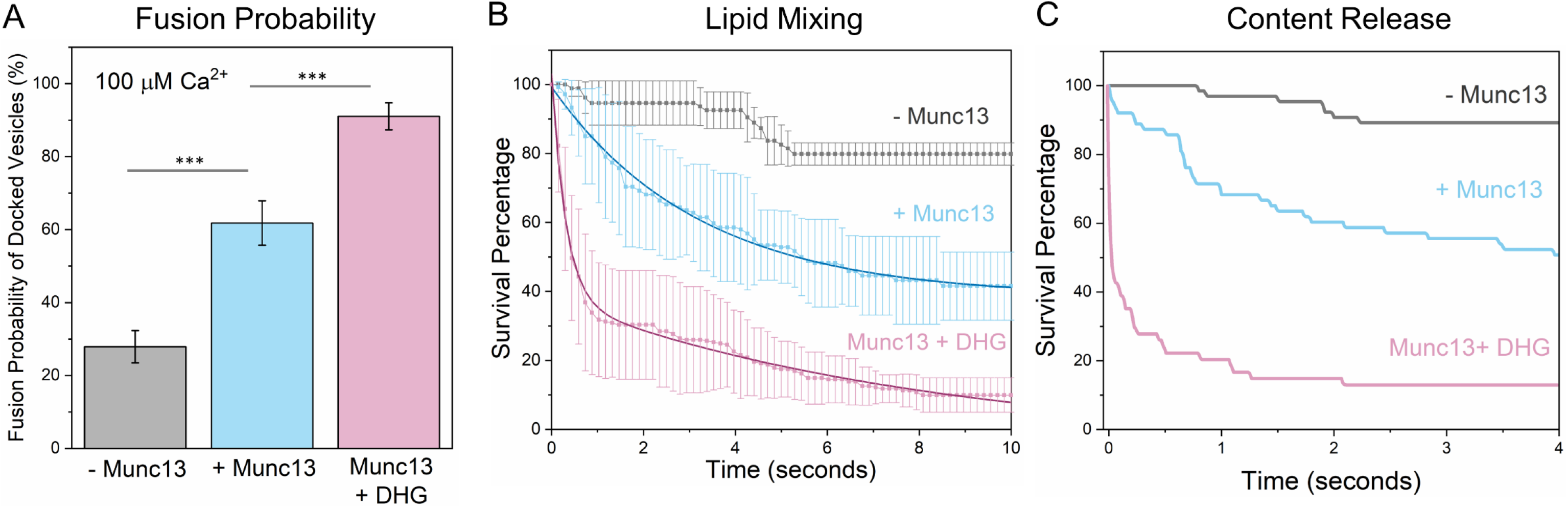
DHG activation of Munc13 is required for rapid, Ca^2+^-evoked vesicular release. The effect of Munc13 and DHG on Ca^2+^ (100 mM)-triggered fusion was evaluated using both lipid mixing (ATTO647N) and content-release (Sulforhodamine-B) assay. Calcium Green, a Ca^2+^-sensor dye, introduced in the suspended bilayer (*via* a lipophilic 24-carbon alkyl chain) was used to monitor the arrival of Ca^2+^ at or near the docked vesicles. (A) End-point analysis (from lipid mixing assay) at 1 min post-Ca^2+^-addition revealed that in the absence of Munc13 or DHG (grey bar), only ∼25% of stably-clamped immobile vesicles fused following Ca^2+^ addition as compared to ∼60% when Munc13 was included (blue bar). Activation of Munc13 by DHG (pink bar) further enhanced the fusion probability with ∼90% of docked, immobile vesicle fusing in response to the Ca^2+^ signal. Kinetic analyses using lipid mixing (B), or content release (C) shows that Munc13 and DHG also enhance the kinetics of Ca^2+^-evoked fusion. The vesicles clamped in the presence of Munc13 and DHG fused rapidly following Ca^2+^-addition (at t = 0), with the majority fusing <1 sec, compared to slower fusion kinetics observed in the absence of DHG and/or Munc13. In the absence of DHG, the average survival trace was best fitted using a one-phase decay nonlinear regression analysis (solid blue line) which yielded a half-life (t_1/2_) ∼2.2 sec. In the presence of DHG, the survival traces were best approximated using a two-phase decay nonlinear regression analysis (solid red line), which yielded a fast t_1/2_ ∼200 msec and a slow t_1/2_ ∼5.7 sec. This data suggests that DHG activation of Munc13 is essential for ultra-fast Ca^2+^-regulated vesicle fusion. (A, B) The average values and standard deviations from three independent experiments (with ∼200 vesicles per condition) are shown. ** p<0.05, ***p<0.005 using the student’s t-test. Note: The fusion kinetics of ∼50 Sulforhodamine-B loaded vesicles per condition are shown in (C). The small number of vesicles precluded detailed statistical analysis. The non-linear nonlinear regression was performed using the built-in nonlinear regression functions with least squares regression in Prism (GraphPad Software Inc.).

### DAG activation of Munc13 is required to achieve fast, Ca^2+^-coupled vesicle fusion

Since DAG and phorbol esters are known to activate Munc13-1 by binding to its C1 domain (32, 37), we investigated the effect of DAG on Munc13-mediated vesicle docking and fusion. The addition of 1% DAG to the lipid bilayer increased the number of docked vesicles, but most of them (∼90%) fused spontaneously (SI Appendix, Figure S3). This significantly reduced the number of docked immobile vesicles, thus hindering the study of Ca^2+^-evoked fusion characteristics. We observed similar behavior with as little as 0.1% DAG in the bilayer (SI Appendix, Figure S3). We have observed that DAG forms separate domains within the bilayer on which Munc13 clusters, irrespective of the bulk concentration (45-48). This limits the ability to effectively vary the local concentration of DAG.

To obviate these limitations, we utilized a short-chain, water-soluble DAG 1,2-hexanoyl-sn-glycerol (DHG) which allowed us to rigorously control both its concentration and the order of addition. To determine the optimal DHG concentration, we incubated the bilayer with various concentrations of DHG (ranging from 0.25 to 1 μM) for 5 minutes before adding Munc13 (along with SUVs and CPX) and then evaluated its effect on vesicle docking and fusion (SI Appendix, Figure S4). We observed minimal effects of DHG on vesicle docking and fusion at concentrations below 500 nM, while higher concentrations (750 nM or 1 μM) increased the probability of spontaneous fusion events without significantly altering the overall fusion probability, essentially recapitulating the results with long-chain DAG (SI Appendix, Figure S4). We chose to use 500 nM of DHG in our experiments for detailed analysis.

The addition of 500 nM DHG to activate Munc13 increased the number of docked vesicles by ∼40% (Figure 1B). Notably, most of these vesicles (∼75%) remained stably docked in an immobile clamped state (Figure 1C) and reached this state in about 250 msec post-docking (Figure 1D). Upon the addition of Ca^2+^ (100 µM), ∼90% of these docked vesicles fused within 8 seconds (Figure 2). Within this envelope, there were two distinct populations of vesicles. Approximately 70% fused with a half-life (t_1/2_) of 200 msec, while the remaining ∼30% fused (lipid mixing) much more slowly (t_1/2_ ∼5.7 secs) (Figure 2B). We also tested and confirmed these findings with a content-release assay using a smaller set of Sulforhodamine B-loaded SUVs under similar experimental conditions (Figure 2C, SI Appendix, Figure S5) but with a faster video frame rate (13 msec). Control experiments without Munc13 showed DHG alone does not change the vesicle docking, priming and fusion characteristics (SI Appendix, Figure S6). Overall, our findings indicate that Munc13 activation by DAG significantly enhances the formation of stably-docked RRP-like vesicles, and is essential for achieving efficient and fast, Ca^2+^-evoked fusion of these docked vesicles.

### Munc13 promotes and accelerates the SNARE complex assembly

We then investigated whether Munc13’s ability to enhance vesicle docking and fusion is related to its SNARE chaperone function. We utilized the binding of fluorescently-labelled CPX, which only binds to pre-assembled SNAREpins with a 1:1 stoichiometry (10, 42, 49), under TIRF conditions to monitor the efficiency and speed of SNARE complex formation under docked vesicles for different reconstitution conditions (Figure 3).

**Figure 3.**
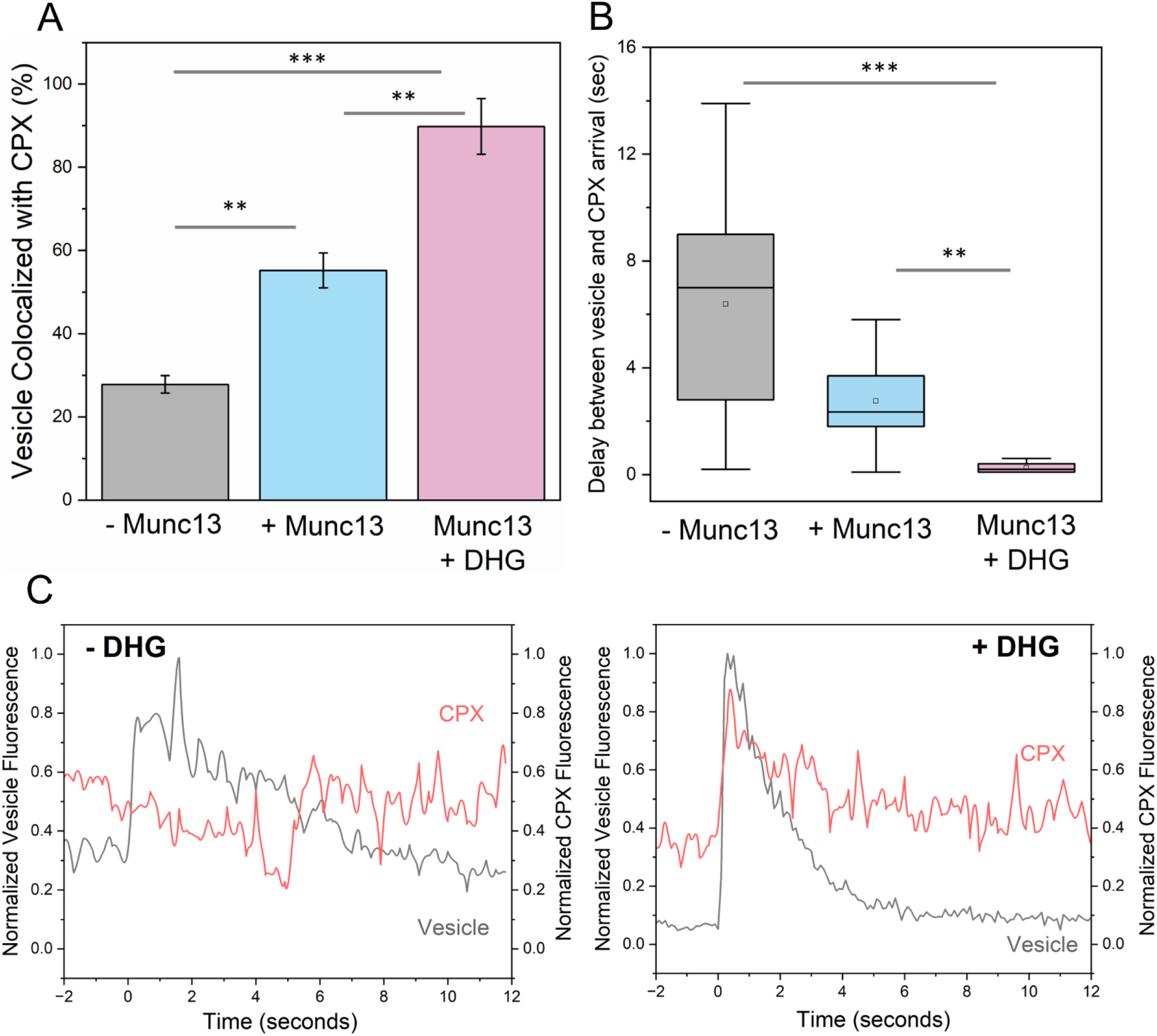
Munc13 and DHG promote and accelerates SNARE complex formation under docked vesicles. The formation of SNAREpins under the docked vesicles (labelled with ATTO465) was visualized by the binding of Alexa Fluor 647-labelled Complexin (CPX). (A) Co-localization analysis revealed that in the absence of Munc13 and DHG (grey bar), only about 25% of vesicles contained CPX signals. The proportion of vesicles colocalized with CPX improved to ∼55% when Munc13 was included (blue bar) and to ∼90% with DHG activation. This indicated that the Munc13 directly promotes the SNARE complex formation, and its chaperone function is further stimulated by DHG binding. (B) Munc13 and DHG also significantly reduced the delay between vesicle docking and CPX binding, with near-simultaneous CPX arrival (<0.2 sec) with DHG activation, compared to a 2-3 sec delay with Munc13 alone and an 8-10 sec delay without Munc13 or DHG. (C) Representative fluorescence traces of vesicle (black) docking and CPX (red) arrival in the presence of Munc13 without or with DHG are shown. The average values and standard deviations from three independent experiments (with ∼100 vesicles per condition) are shown. ** p<0.05, ***p<0.005 using the student’s t-test.

In the absence of Munc13 or DHG, approximately 25% of docked vesicles colocalized with CPX. We also observed significant and variable delays between vesicle docking and the arrival of CPX molecules, with an average delay of around 6 seconds (Figure 3). When Munc13 was present, the delay was reduced to about 3 seconds, and around 55% of vesicles colocalized with CPX. DHG addition greatly improved the efficiency of CPX binding (now ∼ 90% of docked vesicles contained CPX) and greatly reduced the delay in CPX binding (delay of approximately 300 msec). As a control, we tested and verified that CPX mutations (R48A Y52A K69A Y70A; CPX^4A^) that disrupt its interaction with SNAREpins completely abolished its colocalization with the docked vesicles (SI Appendix Figure S7). Our data demonstrate that Munc13 promotes and accelerates the formation of SNAREpins under ready-release vesicles and the Munc13-chaperoned SNAREpin assembly is dramatically accelerated (>10-fold) by DAG binding. Notably, the CPX colocalization levels observed under different reconstitution conditions (Figure 3) are correlated with the fusion probability (Figure 2), suggesting that the efficiency of SNAREpin formation i.e., SNARE priming could account for the observed fusion characteristics.

### Validation of the Munc13/DHG dependent reconstitution

Previous *in vitro* reconstitutions (17, 50) have shown DAG to stimulate Munc13 -dependent Ca^2+^-triggered vesicle fusion, but its necessity was not established. Therefore, it was important to establish the physiological relevance of our DHG-dependent system. To this end, we used the optimized reconstitution protocol (with 500 nM DHG) and a comprehensive suite of mutations whose mechanism of action with respect to Munc13 activation of SNARE assembly has been established (26, 30, 44, 51). This included Munc13 mutations that disrupt Syntaxin1 activation (N1128A/F1131A; Munc13^NFAA^) and VAMP2 recruitment (D1358K; Munc13^DK^) (21, 24, 30); as well as a novel SNAP25 variant (SNAP25^G4S^), in which the last 25 residues (residue 113-138) in the linker region were replaced with a flexible, non-specific GGGGS repeat sequence, rendering it unable to bind to Munc13 (SI Appendix, Figure S8).

None of the SNARE-binding mutants affected the total number of vesicles that attach during the observation (3 min) period (Figure 4A). However, now a vast majority of initially docked vesicles eventually undock (∼75%) or fuse spontaneously (∼5%) during the initial observation period, with only a minor fraction (∼20%) remaining stably docked (Figure 4B). A further 20% of these immobile, ‘clamped’ vesicles fused upon adding 100 µM Ca^2+^, representing only ∼5% of the originally docked vesicles. Similar behavior was observed (Figure 4) with a complementary Syntaxin mutation (R151A/I155A, Syntaxin^RIAA^) that abrogates its interaction with Munc13 (30). These mutations resulted in outcomes indistinguishable from omitting Munc13 protein altogether (Figures 1 and 2) implying that the same binding sites on Munc13 required for neurotransmitter release *in vivo* and SNARE assembly *in vitro* are also required for the priming of ready-release vesicles in the fully defined reconstituted system.

**Figure 4.**
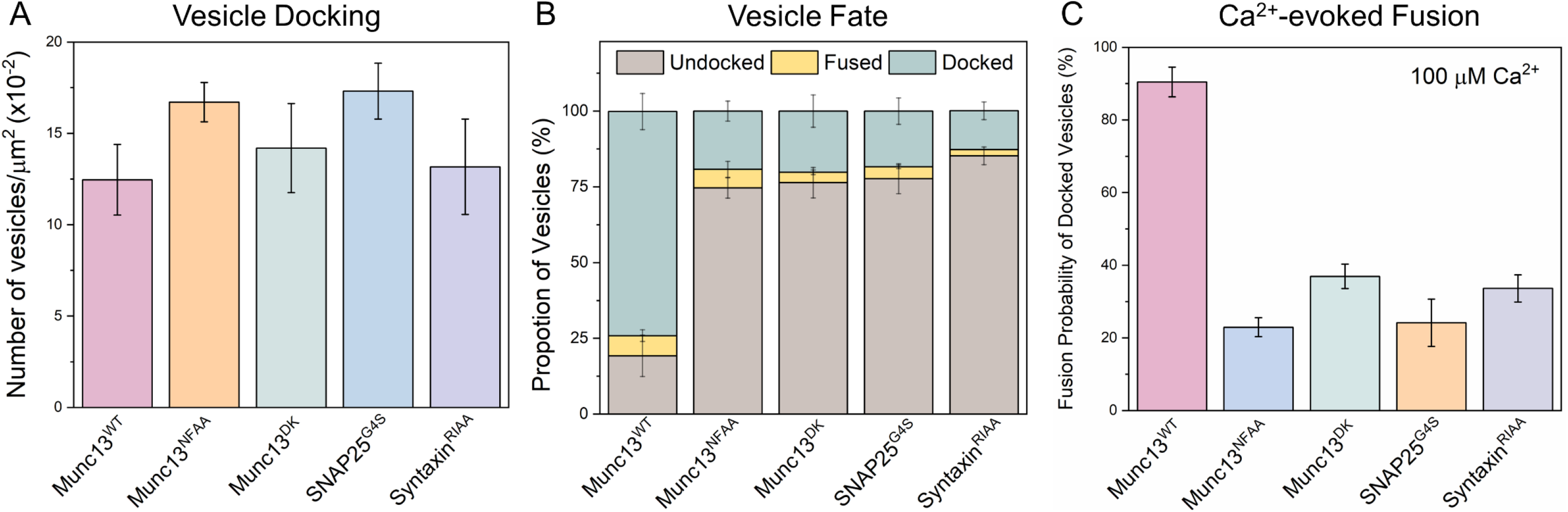
Munc13 interaction with all three SNARE proteins is required for its chaperone function. We employed targeted mutations in Munc13 and SNARE proteins to investigate the functional significance of the Munc13-SNARE interaction in DHG-activated conditions. We used established mutations in Munc13 that disrupt its interaction with Syntaxin (Munc13^NFAA^, orange) or VAMP2 (Munc13^DK^, green) and a reciprocal Syntaxin variant (Syntaxin^RIAA^, purple). Additionally, we designed and tested a novel SNAP25 linker mutant (SNAP25^G4S^, blue) that is incapable of binding Munc13 (SI Appendix, Figure S6). (A) The total number of docked vesicles was similar across all conditions, indicating that the mutations did not affect the initial attachment of vesicles, i.e., docking. (B) Disrupting Munc13’s interaction with any of the SNARE proteins caused most of the vesicles to ultimately dissociate from the bilayer (undocked). (C) The remaining few immobile (“docked”) vesicles failed to fuse upon Ca^2+^ (100 μM) addition The average values and standard deviations from three independent experiments (with ∼200 vesicles per condition) are shown.

### Munc18 and Munc13 cooperative to facilitate priming of reconstituted ready-release vesicles

We also assessed the importance of the Munc18-Syntaxin-VAMP2 ‘template’ complex (23, 25, 52) in the Munc13/DHG-regulated vesicle priming and fusion process (Figure 5). We found that mutations in Munc18 (L348R, Munc18^LR^) or VAMP2 (F77E, VAMP2^FE^), which destabilize the template complex (26, 52), significantly reduced the number of immobile docked vesicles (from ∼75% to ∼25%) and the probability of Ca^2+^-evoked fusion (from ∼85% to ∼20%) of the docked vesicles (Figure 5). Complementary mutations in Munc18 (D326K, Munc18^DK^) that promote or stabilize the template complex (26, 51) also impaired the formation of the immobile docked state but did not affect their fusogenicity (Figure 5). In particular, the Munc18^DK^ reduced undocking (from ∼70% to ∼33%) but strongly increased spontaneous fusion (increasing from ∼5% to ∼25%). As a result, the proportion of immobile docked vesicles was reduced to ∼40%, but the vast majority of these docked vesicles (∼85%) still fused upon Ca^2+^ addition indicating that they had sufficient SNARE assembly. Notably, the omission of Munc13 starkly changed the probability and kinetics of Ca^2+^-evoked fusion involving the Munc18^DK^ mutant (SI Appendix, Figure S9). In the presence of Munc13, Ca^2+^-coupled release occurred with >90% probability and within 2 secs. In the absence of Munc13, only 50% of docked vesicles fused and it extended >10 sec post-Ca^2+^ addition (SI Appendix, Figure 9). As expected, none of the mutants tested affected the number of docked vesicles (Figure 5). Taken together, our data indicate that the Munc18-Syntaxin-VAMP2 ‘template’ complex is a key functional intermediate in the SNARE assembly pathway and is required for the formation of ready-release vesicles and for their subsequent Ca^2+^-evoked fusion in the fully reconstituted system and coordinated action of both Munc13 and Munc18 is required.

**Figure 5.**
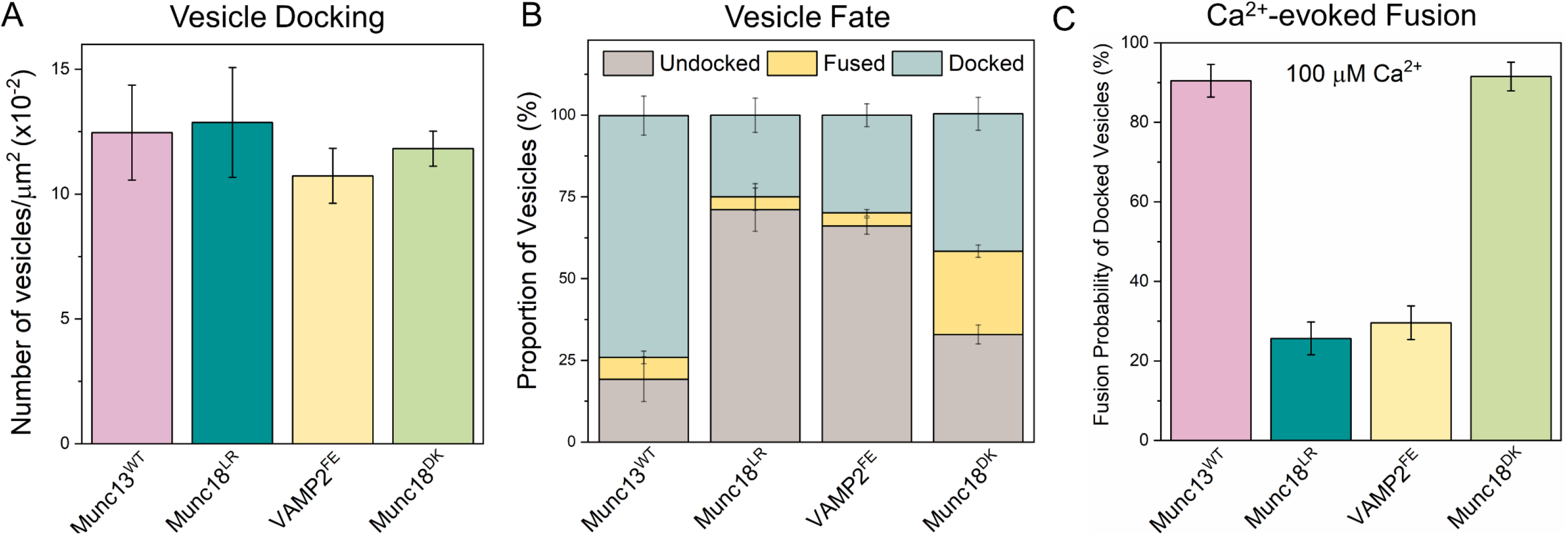
Munc18 and Munc13 act cooperatively to facilitate SNARE complex assembly via the ‘template’ complex. To investigate the role of the Munc18-Syntaxin-VAMP2 ‘template’ complex in the Munc13-mediated vesicle priming and fusion process, we examined the effects of mutations that either disrupt (Munc18^LR^, teal; VAMP2^FE^, yellow) or promote (Munc18^DK^, green) template complex formation. (A) We observed a similar number of vesicles for all mutations tested, suggesting that template complex formation is not essential for vesicle docking. (B) The fate of the docked vesicles, however, differed depending on the mutation. Negative mutations (Munc18^LR^ and VAMP2^FE^) led to vesicle undocking, while the positive mutation (Munc18^DK^) potentiated spontaneous fusion. Overall, both the positive and negative mutations decreased the number of docked ‘immobile’ vesicles. (C) Disrupting template complex formation also abolished the Ca^2+^-evoked fusion of docked vesicles. Taken together, the data indicate that template complex formation is necessary for productive SNARE complex formation and the coordinated action of Munc13 and Munc18 is required to generate stably docked vesicles. The average values and standard deviations from three independent experiments (with ∼200 vesicles per condition) are shown.

## DISCUSSION

Here we introduce a novel biochemically-defined fusion system that provides the most physiologically accurate *in vitro* reconstitution of synaptic fusion machinery to date. The system includes the first reconstitution of the functional Syntaxin:Munc18 complex along with palmitoylated SNAP25 into a pore-spanning lipid bilayer mimicking pre-synaptic membranes. This represents a significant improvement over previous efforts which involved either pre-assembled t-SNAREs (bypassing the need for both Munc18 and Munc13) or soluble Munc18 and SNAP25 added in large excess (13, 17, 39, 42, 50). These advances were facilitated by the development of optimized protocols enabling us to routinely purify a stable 1:1 Syntaxin:Munc18 complex using Triton X-100 detergent and *in vitro* palmitoylation of SNAP25 (see Materials & Methods for details). Combining this bilayer system with SUVs mimicking synaptic vesicles containing physiologically-relevant numbers of VAMP2 and Synaptotagmin allowed us to dissect the interplay between Munc18, Munc13 and DAG in vesicle docking, priming, and Ca^2+^-evoked fusion in molecular terms.

We find that DHG acts as an essential co-factor and is required for optimal Munc13 chaperone function (Figures 1 & 2). The activation of Munc13 by DAG significantly improved the formation of stably docked RRP-like vesicles and is essential for achieving high-efficiency Ca^2+^-evoked release. However, the effect of short-chain DAG was concentration-dependent - it facilitated Ca^2+^-evoked response at low concentrations but potentiated both spontaneous and evoked fusion at higher concentrations. The behavior observed under high DHG concentration closely resembles the effect of phorbol ester on neurotransmitter release in cultured neurons (53).

We observed that the inclusion of long-chain DAG (even at 0.1%) in the bilayer resulted in extensive spontaneous vesicle fusion, so we used DHG, a soluble DAG analogue, in our experiments. DHG has the same headgroup as DAG and thus, is expected to bind to the Munc13 C1 domain similarly with comparable affinity. Indeed, adding DHG in solution (500 nM) promoted Munc13 to bind to lipid bilayers and to a similar extent as including 1% long-chain DAG in the bilayer (SI Appendix, Figure S10A). DHG, similar to DAG, promoted the formation of Munc13 clusters on the membrane surface (SI Appendix, Figure S10B) further arguing that DHG is a suitable substitute for DAG. We suspect that the formation of DAG-rich domains, which are likely to separate from the fluid phospholipid bilayers (45-48), accounts for the discrepancy between DAG and DHG in our experimental condition.

This phenomenon is likely created by what amounts to an *in vitro* artefact because such DAG islands are unlikely to exist in neuronal active zones where the abundance of DAG is dynamically and spatially limited and occurs in the context of PIP_2_-rich nanodomains (54, 55). DAG is produced locally by the hydrolysis of PIP_2_ by Ca^2+^-regulated phospholipase C, and the excess DAG is rapidly converted to phosphatidic acid by diacylglycerol kinase and ultimately recycled to PIP_2_ (55, 56). The coordinated action of these enzymes likely ensures that DAG is presented to Munc13, and other active zone proteins as a molecularly dilute and dispersed component mixed within the ∼ 70 nm diameter PIP_2_-rich nanodomains that are known to assemble cluster around Syntaxin both *in vivo* and in reconstituted bilayers (57). Dynamic alterations of DAG levels in response to Ca^2+^ signals have been shown to play a role in short-term plasticity. Specifically, the activation of the Munc13 C_1_ domain by DAG resulting from Ca^2+^-dependent activation of phospholipase C has been shown to potentiate Munc13 function, resulting in short-term synaptic facilitation (38, 58).

By using fluorescent CPX binding as a label for SNAREpin formation (Figure 3), we confirmed that the ability of Munc13 to stimulate vesicle fusion probability and kinetics is directly linked to its SNARE chaperone function. Specifically, we find that the improved fusion characteristics observed with Munc13 ± DAG are directly correlated with the enhanced efficacy of SNARE complex formation. This suggested that the SNARE priming process is the rate-limiting step in the vesicle fusion pathway. Supporting this proposition, we have observed a large proportion of stably docked immobile vesicles with very little to no undocking when co-purified t-SNARE complex is used (13, 39, 42), as it is expected to enhance the rate of productive ternary SNARE complex formation.

Systematic mutational analyses confirmed that Munc13 and Munc18 act cooperatively, with intersecting interactions with individual SNARE proteins, to guide efficient SNARE complex assembly. Munc18 binds to Syntaxin and VAMP2 to create an intermediate Munc13-Syntaxin-VAMP2 ‘template’ complex that is critical for productive SNARE complex formation and vesicle fusion. Munc13’s chaperone function involves interactions with all three individual SNARE proteins. It accelerates and stabilizes the ‘template’ complex by promoting proper alignment of Syntaxin and VAMP2. Additionally, Munc13 recruits SNAP25 to the ‘template’ complex, stimulating ternary SNARE complex formation.

Our data further suggest that the activities of Munc18 and Munc13 are closely coordinated and must be balanced for effective clamping of SNAREpins to produce stably docked vesicles and to elicit fast Ca^2+^-evoked vesicular fusion. This implies that SNARE assembly occurs in stages, with sequential and overlapping interactions with chaperones (Munc18 and Munc13) and regulatory elements (Synaptotagmin and Complexin). This ensures that a specific number of SNAREpins are produced and then firmly held in a release-ready state, to facilitate cooperative zippering and ultra-fast release upon Ca^2+^ influx. How this occurs in molecular terms is not known.

Insight into the possible mechanism can be gleaned from recent cryoEM structural analyses which revealed that Munc13 undergoes distinct structural transitions involving different oligomeric states (27). We posit that these topological transitions, which are implied by the structures observed to be intimately linked to DAG and Ca^2+^-binding, likely choreograph the SV docking/priming process (27). As such, Munc13 functions as a molecular scaffold that helps to organize the exocytic proteins into a functional complex that is necessary for efficient vesicular release. The *in vitro* fusion system we describe here provides a unique, focused, and powerful tool to test this specific and other molecular models, and progress towards the goal of deciphering the pointillistic details of SNARE priming by Munc18 and Munc13.

## MATERIALS AND METHODS

### Materials

The following cDNA constructs previously described (13, 39, 44) were used in this study: full-length VAMP2 (mouse His^6^-SUMO-VAMP2-, residues 1-116); full-length SNAP25 (mouse His^6^-SNAP25b, residue 1-206); Synaptotagmin (rat Synaptotagmin1-His^5^, residues 57-421); Complexin (human His^6^-Complexin 1, residues 1-134) and Munc13 C1-C2B-MUN-C2C domain (rat His^12^-Munc13-1, residues 529-1735 with residues 1408–1452 replaced by the sequence EF and 1532-1550 deleted). We generated a new expression clone in the pET28vector to express and purify the Munc18/Syntaxin complex (rat His^6^-SUMO Munc18/Syntaxin1). Phusion High Fidelity Mastermix (New England Biolabs, Ipswich, MA) was used to generate variants in Munc13 (Munc13^NFAA, DK^) VAMP2 (VAMP2^FE^), Syntaxin (Syntaxin^RIAA^) and Munc18 (Munc18^LR, DK^). The SNAP25 linker mutants (SNAP25^G4S^) were synthesized (Genewiz, Inc., Plainfield, NJ) with the SNAP25 loop residues 82–105, 97-126 and 113-138 replaced with (GGGGS) repeats and cloned into the pET28 vector similar to the SNAP25^WT^ construct.

Lipids including 1, 2-Dioleoyl-sn-glycero-3-phosphocholine (DOPC), 1, 2-Dioleoyl-sn-glycero-3-phospho-L-serine (DOPS), L-α-phosphatidylinositol-4, 5-bisphosphate (Brain PIP2) and 1, 2-Dioctadecanoyl-sn-glycerol (DAG) were purchased from Avanti Polar Lipids (Alabaster, AL). Fluorescent lipids ATTO465-DOPE and ATTO647N-DOPE were purchased from ATTO Tec (Siegen, Germany). 1, 2-Dihexanoyl-sn-glycerol (DHG) was purchased from Cayman Chemicals (Ann Arbor, MI). Alexa Fluor 568/647 – Maleimide and TCEP HCl were purchased from Thermo Fisher (Waltham, MA). Calcium Green conjugated to a lipophilic, 24-carbon alkyl chain (Calcium Green C24) was custom synthesized by Marker Gene Technologies (Eugene, OR).

### Protein Expression and Purification

All SNARE and associated proteins were expressed and purified as described previously (13, 39, 44). Briefly, the proteins were expressed in Escherichia coli strain BL21(DE3) (Novagen, Darmstadt, Germany) and cells were lysed with a cell disruptor (Avestin, Ottawa, Canada) in lysis buffer containing 50 mm HEPES, pH 7.4, 400 mM KCl, 2% Triton-X 100, 10% glycerol, 1 mm Tris (2-carboxyethyl) phosphine hydrochloride (TCEP), and 1 mm phenylmethylsulfonyl fluoride. Samples were clarified using a 45 Ti rotor (Beckman Coulter, Brea, CA, USA) at 140, 000 x *g* for 30 minutes and incubated with Ni-NTA agarose (Thermo Fisher, Waltham, MA, USA) for 4– 16 hours at 4 °C. The resin was subsequently washed (3 column volumes) in the wash buffer (50 mM HEPES, 400 mM KCl, 10% Glycerol, 1mM TCEP, 32 mm imidazole) with no detergent (Complexin and SNAP25) or 1% Octylglucoside (VAMP2 and Synaptotagmin) or 1% Triton X-100 (Munc18/Syntaxin). Note: We carried out systematic detergent (Octylglucoside, Sodium Cholate, CHAPS, Dodecyl Phosphocholine, Triton X-100) screening to identify 1% Triton X-100 as the ideal detergent to produce stable and functional Munc18/Syntaxin complex. The proteins were either eluted with 300 mM Imidazole (Synaptotagmin) or cleaved off the resin with either Thrombin (SNAP25 and Complexin) or SUMO protease (VAMP2 and Munc18/Syntaxin) in HEPES buffer for 2 hours at room temperature. SNAP25 and Complexin proteins were further purified using gel filtration (Superdex75 Hi-load column, GE Healthcare, Chicago, IL), and Synaptotagmin-1 protein was subjected to anion exchange chromatography (MonoS, GE Healthcare, Chicago, IL) to remove nucleotide contaminants. The peak fractions were pooled and concentrated using filters of appropriate cutoffs (EMD Millipore, Burlington, MA, USA). SNAP25 was then palmitoylated using a 20-fold excess of Palmitoyl Coenzyme A (Sigma Aldrich, St Louis, MO) in HEPES buffer supplemented with 1% TritonX-100 for 30 minutes at room temperature with gentle mixing. Native mass spectrometry analysis confirmed the palmitoylation of SNAP25 molecule (SI Appendix, Figure S11A) and combined with float-up analysis, we estimate 30-50% of proteins are palmitoylated.

Munc13 was expressed in ExpiHEK-293 cell cultures using ExpiFectamine as a transfection reagent (Thermo Fisher, Waltham, MA). Briefly, thawed cells were passaged three times prior to transfection and were grown for 72 hours before being spun down and rinsed in ice-cold PBS. The pellet was resuspended in lysis buffer and lysed using a Dounce homogenizer. The sample was clarified at 140, 000 *g* for 30 minutes at 4 °C and the supernatant was incubated overnight with Ni-NTA beads, in the presence of DNAse1, RNAseA, and Benzonase to remove nucleotide contamination. The protein was further washed in the lysis buffer (without Triton-X 100) before being cleaved with PreScission protease for 2 hours at room temperature. The eluted proteins were further purified via gel filtration (Superdex 200, GE Healthcare Chicago, IL, USA). In all cases, the protein concentration was determined using a Bradford Assay (Bio-Rad, Hercules, CA, USA), with BSA as a standard, and protein purity was verified using SDS/PAGE analysis with Coomassie stain. All proteins were flash-frozen and stored at −80 °C for long-term storage.

### Suspended Bilayer and Vesicle Preparation

For the suspended bilayer, we approximated the pre-synaptic membrane physiological composition with 81% DOPC, 15% DOPS, 3% PIP2 and 1% ATTO465-PE for visualization. The lipids were mixed and dried under N_2_ gas followed by a vacuum. For a subset of experiments, we included Ca^2+^-sensor, Calcium-green containing a 24-carbon alkyl chain during the sample preparation. Bilayer samples were rehydrated with Munc18/Syntaxin1 and palmitoylated SNAP25 (1:3200 protein: lipid input ratio) in 5x buffer (125 mM HEPES, 600 mM KCl, 1 mM TCEP, pH 7.4) supplemented with 2% TritonX-100 for thirty minutes. Samples were then mixed directly with SM-2 Biobeads (Bio-Rad, Hercules, CA) for 30 minutes with gentle agitation. The samples were further dialyzed overnight in 5X buffer without detergent. We used mass spectrometry to verify the complete removal of Triton X-100 (SI Appendix, Figure S11B).

Lipid bilayers were created by drying and rehydrating membranes to form GUVs as previously described (40). Briefly, 4 μL drops of Munc18/Syntaxin/SNAP25 containing proteoliposomes were dried on a clean Mattek dish and rehydrated twice. In the second rehydration, the sample was diluted 5X to 20 μL with distilled water and then added to a cleaned silicon chip containing 1x buffer (25 mM HEPES, 120 mM KCl, 1 mM TCEP, pH 7.4) supplemented with Mg^2+^ (5 mM). The bilayer was extensively washed with 1x buffer, and the fluidity of the suspended bilayer was verified using fluorescence recovery after photobleaching (FRAP) using ATTO465 (lipid) or Alexa568 (Protein) fluorescence labels. Consistent with a fluid and mobile bilayer, the average diffusion coefficient of the lipid was calculated to be 3.8 μm^2^/s while the protein diffusion coefficient was found to be 1.7 μm^2^/s (SI Appendix, Figure S12). For TIRF experiments, we created the suspended bilayer using the redesigned silicon chip (41), with buffer supplemented with 45% OptiPrep gradient media for index matching (41). Optiprep on the top was washed out with 1X buffer after bilayer formation. Bilayers contained 0.1% ATTO465 and were photobleached with 100% laser power after verifying they had formed.

For small-unilamellar vesicle preparation, we approximated the synaptic vesicle lipid composition using 83% DOPC, 15% DOPS, 2% ATTO647N-PE. The samples were dried under N_2_ gas followed by a vacuum. Lipids were rehydrated with VAMP2 (1:100) and Synaptotagmin1 (1:250) in buffer (140 mM KCl, 50 mM HEPES, 1 mM TCEP, pH 7.4) supplemented with 1% Octyl β-glucoside. After 30 minutes of mixing, samples were rapidly diluted 3X below CMC and allowed to sit for another 30 minutes before being dialyzed overnight buffer without detergent. The samples were subjected to additional purification on the discontinuous Nycodenz gradient. For content-release assays, we prepared vesicles supplemented with 25 mol% cholesterol in the lipid formulation with 30 mM Sulforhodamine-B. Samples were subjected to gel filtration using a CL4B column to remove the unincorporated dye and were further purified as described above. Based on the densitometry analysis of Coomassie-stained SDS gels and chymotrypsin digest, we estimated vesicles contained ∼74 and ∼25 copies of outward-facing VAMP2 and Syt1 respectively

### Single Vesicle Fusion Assays

Single vesicle assays were performed as described previously (13, 42) with a few modifications. DHG (at the defined concentration) was added to the pre-formed bilayer and incubated for 5 minutes. Vesicles (500 nM lipids) were added from the top using a pipette and allowed to interact with the bilayer for 3 minutes. We used the ATTO647-PE or Sulforhodamine-B fluorescence to track the fate of the individual vesicles. All vesicles that attach to the suspended bilayer during the 3-minute observation period were considered as “docked” vesicles. Initially, all docked vesicles exhibited diffusive mobility and subsequently transitioned to an ‘immobile’ clamped state (SI Appendix Figure S2). Among these mobile and immobile vesicles, some underwent spontaneous fusion, evident by a burst of fluorescence intensity followed by a rapid decrease. Furthermore, we observed instances of undocking, where vesicle fluorescence disappeared without a fusion burst, primarily during the mobile phase (SI Appendix Figure S2). Undocking events were infrequent once vesicles reached the immobile state.

After the initial 3 min interaction phase, the excess vesicles in the chamber were removed by buffer exchange (3X buffer wash) and 100 µM CaCl_2_ was added from the top to monitor the effect of Ca^2+^ on the docked vesicles. The lipid mixing experiments were recorded at a 147 msec frame rate to track large numbers (∼30-40 vesicles) simultaneously. For the content-release assay, the frame rate was increased to 13 msec by adjusting to a smaller region of interest and thus, tracking fewer vesicles at a time. As before, the increase in Calcium green fluorescence at 532 nm was used to precisely estimate the arrival of Ca^2+^ at or near the docked vesicle (SI Appendix, Figure S4).

### Complexin Binding Assay

To track the binding of Complexin to docked vesicles, we used the current iteration of the silicon chips which are compatible with TIRF microscopy (41). For these co-localization experiments, vesicles were labelled with 2% ATTO465-PE and the sole Cysteine residue in Complexin1 was labelled with AlexaFluor 647-Maleimide, with >85% efficiency. The bilayer was photobleached prior to the addition of vesicles, mixed with labelled CPX and Munc13, without or with pre-incubation with DHG. We used a DuoView 2 system for simultaneous tracking of vesicles and CPX in the 488 and 647 channels. All experiments were carried out at a 100 msec frame rate on TIRF (Nikon) microscope.

### Microscale Thermophoresis Interaction Analysis

Microscale Thermophoresis (MST) analysis was carried out as described previously (44). Briefly, Halo-tagged Munc13-1 construct was labeled with AlexaFluor 660 and MST analysis was carried out with titration of SNAP25 (1 nM – 125 µM range) into 50 nM of fluorescently labeled Munc13-1 constructs in a premium-coated glass capillary using a Monolith NT.115 (Nanotemper Technologies, San Francisco, CA).

### Cryo-electron Microscopy Analysis

The cryo-electron microscopy analysis of Munc13 binding to lipid membrane was carried out described previously (27). Pre-formed vesicles (DOPC/DOPS/PIP2 in a molar ratio of 14/80/6) were diluted down to lipid concentration of 100 μM and mixed with 0.25 μM Munc13C protein in 1:1 (vol/vol) ratio. Once mixed, the samples were incubated at room temperature for 5 min then kept on ice until freezing. Samples (2.5 μl) were vitrified using a Vitrobot Mark IV (Thermo Fisher Scientific) held at 8°C with 100% humidity. They were applied to freshly glow-discharged 200 mesh Lacey Formvar/carbon grids and grids were blotted for 5 s with blot force -1 and then plunged frozen in liquid ethane cooled by liquid nitrogen. Samples were imaged using Glacios Cryo TEM 200 kV (Thermo Fisher Scientific, Waltham, MA) equipped with a K2 Summit direct electron detector (Gatan, Pleasanton, CA).

### Munc13 Clustering Assay

We adapted the protocol described previously (36, 44) to assess the formation of Munc13 clusters on lipid membrane surface in the absence or presence of DAG/DHG. Liposomes were prepared with following lipid composition (71% DOPC, 25% DOPS, 2% PIP2, ± 2% DAG) using extrusion method with HEPES buffer (50 mM HEPES, 140 mM KCl, 1 mM TCEP, pH 7.4). Lipid bilayers were created by Mg^2+^ (5 mM) induced bursting liposomes in ibidi glass-bottom chambers (ibidi GmbH, Germany) and extensively washed with the HEPES buffer supplemented with EDTA (6 mM). Munc13-Halo-Alexa488 (10 nM) was added to the pre-washed bilayer and incubated for 60 mins. The samples were imaged on a TIRF (Nikon) microscope with a 63x oil objective. For DHG experiments, we added 500 nM DHG is solution along with Munc13.

### Mass Spectrometry Analysis

To carry out mass spectrometry analysis, all samples were buffer exchanged to 200 mM ammonium acetate with Zeba™ spin desalting columns (Thermo Fisher Scientific, Waltham, MA), with an approximate protein concentration of 2-10 µM. Stable electrospray ionization was achieved using in-house nano-emitter capillaries in Q Exactive UHMR (Thermo Fisher Scientific, Waltham, MA). The nano-emitter capillaries were formed by pulling borosilicate glass capillaries (O.D – 1.2mm, I.D – 0.69mm, length – 10cm, Sutter Instruments) using a Flaming/Brown micropipette puller (Model P-1000, Sutter Instruments, Novato, CA). After the tips were formed using this puller, the nano-emitters were coated with gold using rotary pumped coater Q150R Plus (Quorum Technologies, Laughton, East Sussex, UK).

To perform the mass spectrometry-based measurement, the nano-emitter capillary was filled with buffer-exchanged protein samples and installed into Nanospray Felx™ ion source (Thermo Fisher Scientific, Waltham, MA). The MS parameters were optimized as per the samples of analysis. The typical parameters were as follow: spray voltage was in the range of 1.2 -1.5 kV, capillary temperature was 275^0^C, the resolving power of the MS was between 3,125 – 6,250 at m/z of 400, the ultrahigh vacuum pressure of 5.51e^-10^ – 6.68e^-10^ mbar, the in-source trapping range between -50V and -200V. The HCD voltage was optimized for each sample to achieve a good-quality spectra and the range was between 0 to 200V. All the mass spectra were visualized and analyzed with the Xcalibur software.

## ACKNOWLEDGEMENTS

This work was supported by the National Institute of Health (NIH) grant DK027044 (JR/SK) and GM141194 (KG).

## AUTHOR CONTRIBUTION

R.V.K.S., A.C, J.E.R, S.S.K designed research; R.V.K.S., A.C, M.B, K.G, F.L, A.P performed research R.V.K.S., A.C, M.B, K.G, F.L, A.P, S.S.K analyzed data; J.C, S.R, A.E contributed reagents, R.V.K.S., S.S.K, J.E.R wrote the paper. All authors read and revised the manuscript.

