## Supplementary Information for "Novel Roles for Diacylglycerol in Synaptic Vesicle Priming and Release Revealed by Complete Reconstitution of Core Protein Machinery"

### SI Appendix

[#] These authors contributed equally

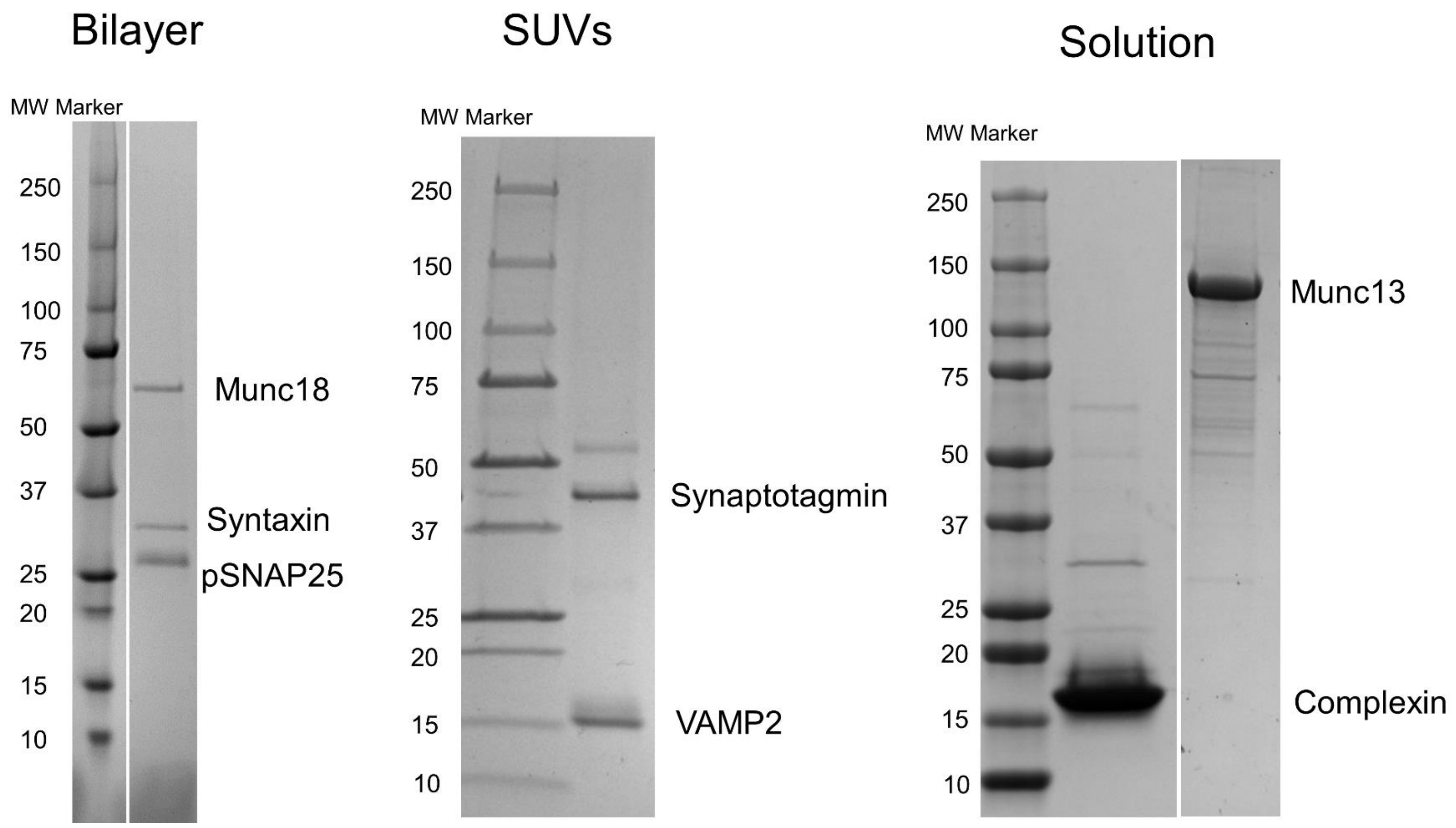


***Figure S1.*** Coomassie gel image of proteins used in this study. The proteins were purified and reconstituted into the suspended bilayer (left) or SUVs (middle) as described in the Methods section.

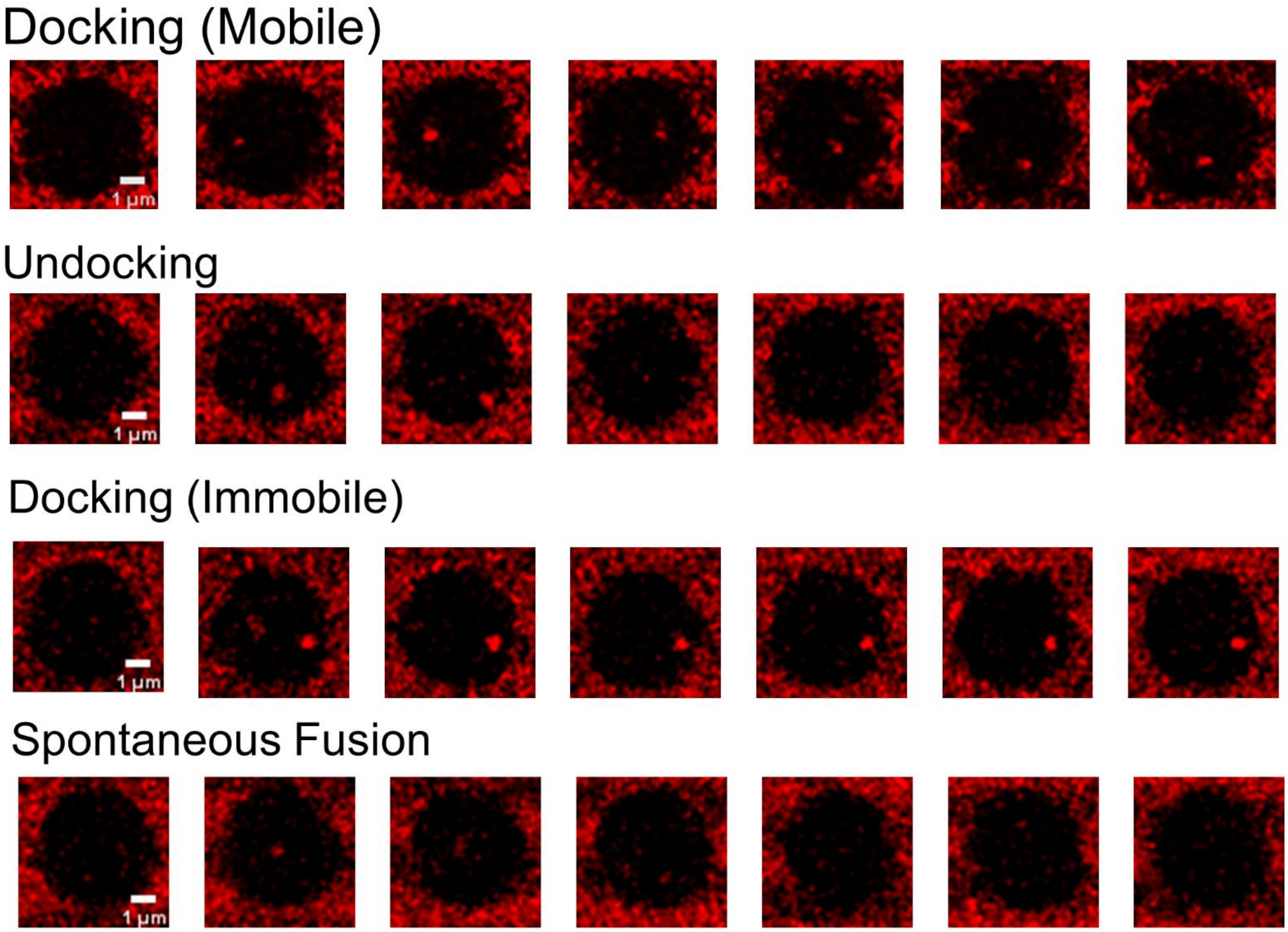


***Figure S2.*** Representative time-lapse fluorescence (ATTO647N) images (frame rate = 147 ms) of single docked SUVs showing the behavior of docked vesicles observed under different molecular conditions. In all cases, the vesicle docks to the bilayer surface at the 2$^{nd}$ frame. The docked vesicles are mobile on the bilayer surface (1$^{st}$ panel) and sometimes undocks (2$^{nd}$ panel) as evidenced by the disappearance of the vesicle fluorescence signal. Some vesicles reach the stable, immobile state (3$^{rd}$ panel) and some of the immobile vesicles spontaneously fuse (4$^{th}$ panel) which is observed by the spreading of the vesicle fluorescence signal.

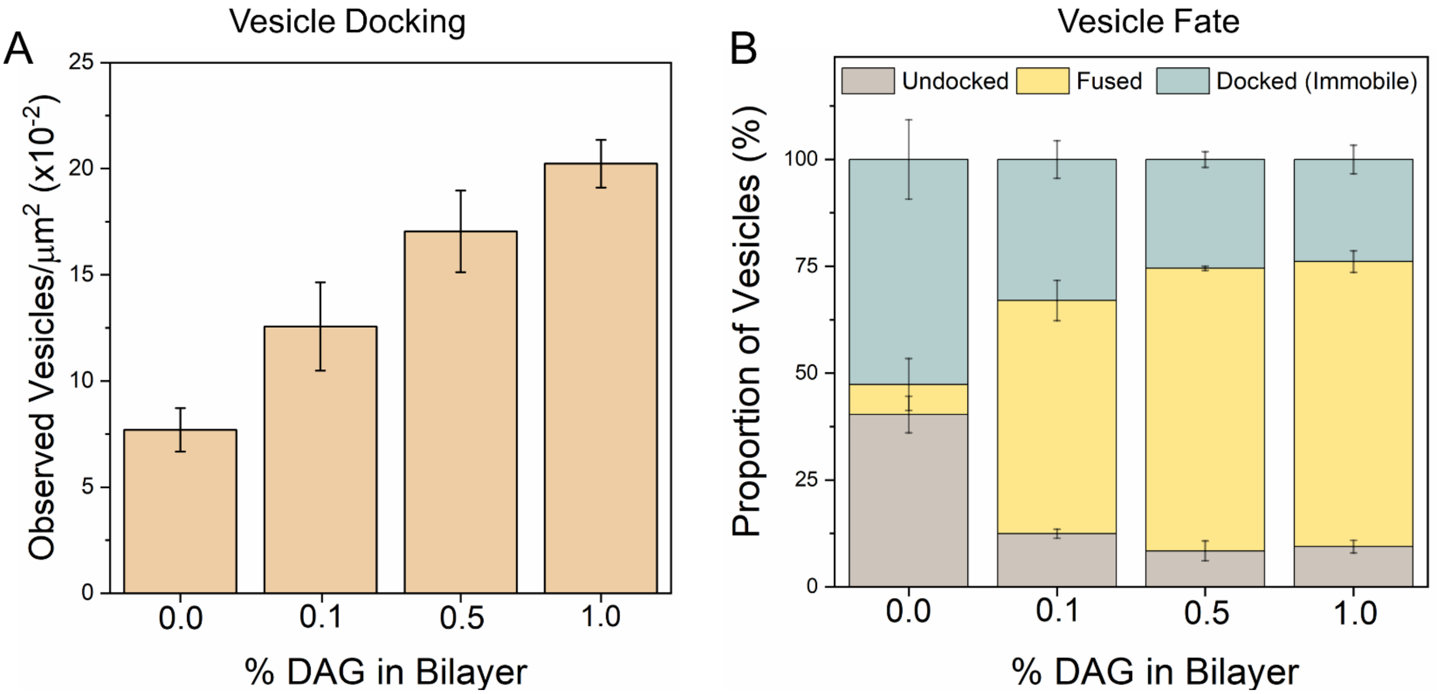


***Figure S3.*** The inclusion of diacylglycerol (DAG) in the bilayer increased the number of docked vesicles (A) but it also potentiated the spontaneous release resulting in a small portion of vesicles converting into the immobile docked state (B). This behavior was observed even at the lowest concentration (0.1%) of DAG tested. The average values and standard deviations from three independent experiments (with ~100 vesicles per condition) are shown.

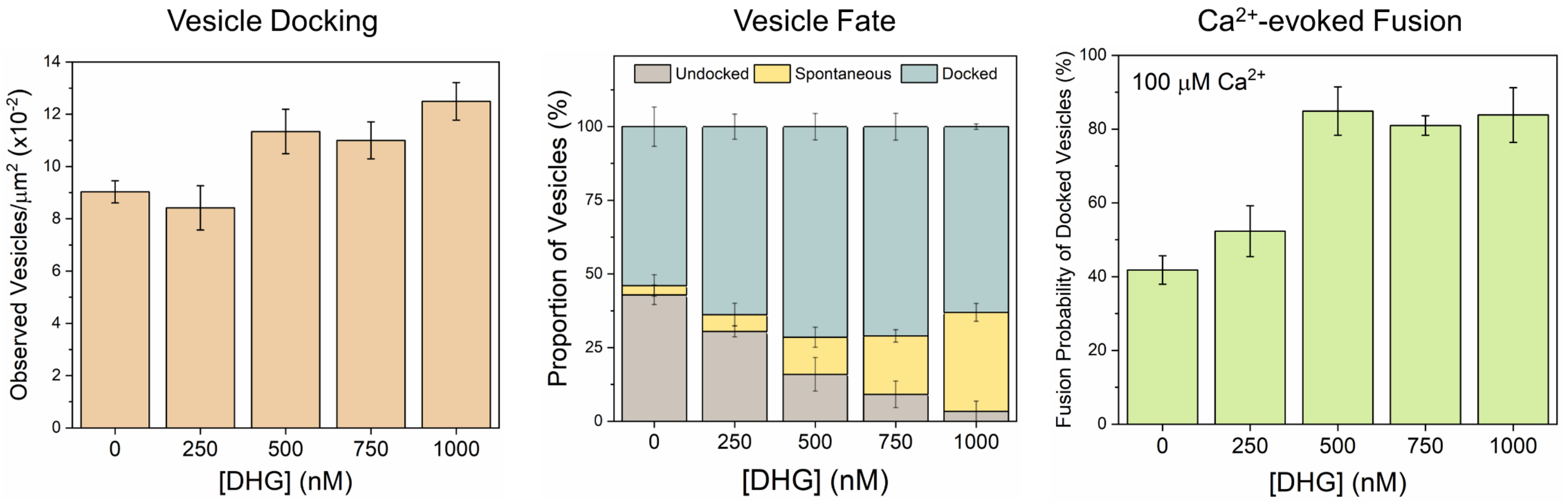


***Figure S4.*** Short-chain DAG (DHG) exhibited a concentration-dependent effect on Munc13-regulated vesicle docking, priming and $Ca^{2+}$-evoked release. DHG at concentrations ≤ 500 nM had very little or no effect. At 500 nM or above, DHG increased the number of vesicles and the $Ca^{2+}$-evoked fusion probability of docked vesicles. DHG also potentiated spontaneous fusion at concentrations ≥750 nM increased, thus lowering the total number of docked immobile vesicles. Hence, we chose to use 500 nM DHG in our experiments. In all experiments, DHG was added in solution 5 min prior to the addition of vesicle/Munc13. The average values and standard deviations from three independent experiments (with ~100 vesicles per condition) are shown.

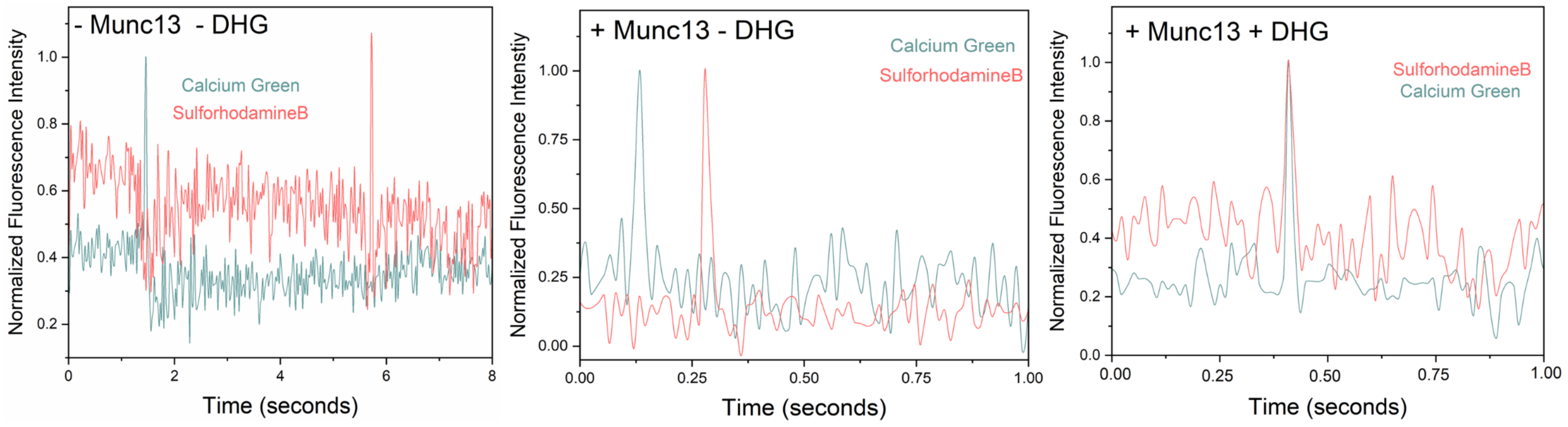


***Figure S5.*** The effect of Munc13 and DHG on $Ca^{2+}$-triggered fusion was assessed using a content-release assay with SulforhodamineB-loaded vesicles. SulforhodamineB is largely self-quenched when encapsulated inside an SUV. Fusion of the vesicle results in dilution of the probe, which is accompanied by increasing fluorescence. The $Ca^{2+}$-sensor dye, Calcium Green, introduced in the suspended bilayer (*via* a lipophilic 24-carbon alkyl chain) was used to monitor the arrival of $Ca^{2+}$ at/near the docked vesicles. Representative fluorescence traces before and after the addition of 100 μM $Ca^{2+}$ under different reconstitution conditions show that the rise in Sulforhodamine-B (red curve) fluorescence intensity occurs within a single frame (13 msec) of $Ca^{2+}$ binding to local Calcium green (green curve) in the presence of Munc13 and DHG (Right panel). The vesicle fusion is slow if DHG is omitted (Middle panel), and even slower when both Munc13 and DHG are omitted (Left panel).

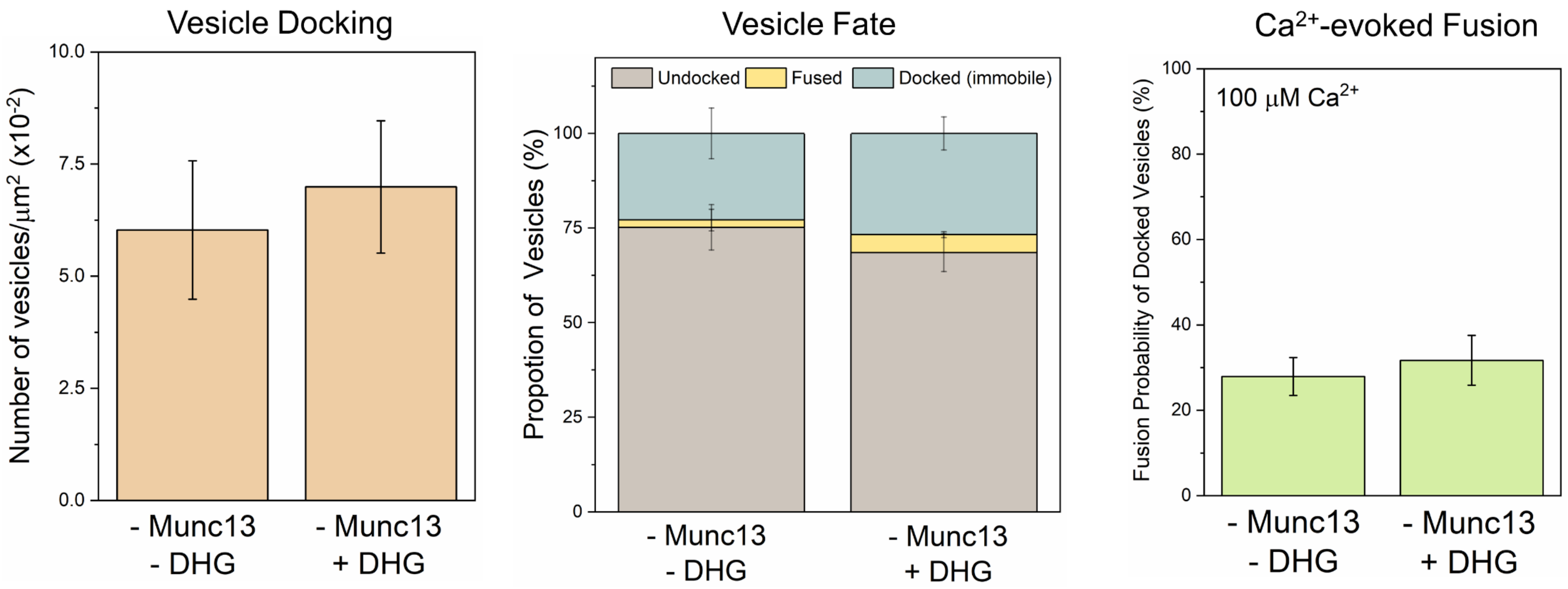


***Figure S6.*** The inclusion of DHG (500 nM) without Munc13 did not change the vesicle docking, priming and $Ca^{2+}$-evoked release characteristics. This data confirms that DHG activation of Munc13 is required to produce stably docked immobile vesicles and elicit efficient fusion in response to $Ca^{2+}$-signal. The average values and standard deviations from three independent experiments (with ~100 vesicles per condition) are shown.

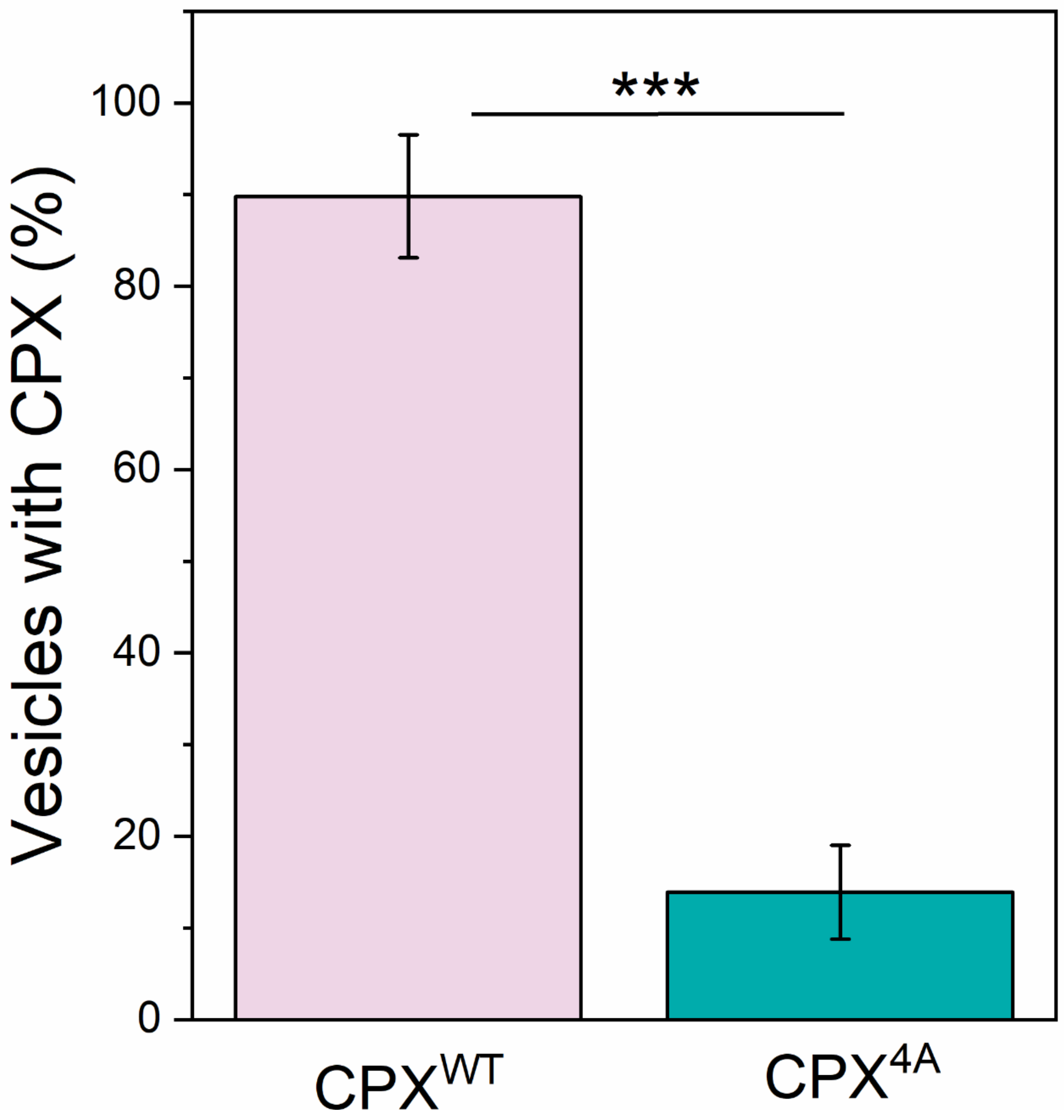


***Figure S7.*** TIRF analysis using Alexa Fluor 647-labelled Complexin (CPX) and ATTO465-labelled vesicles shows that disrupting the interaction of CPX central helix to the SNARE complex using targeted mutations (R48A Y52A K69A Y70A; $CPX^{4A}$) eliminates the colocalization. This indicates the CPX binding and colocalization are well-suited to track the efficiency and speed of SNARE complex formation under docked vesicles.

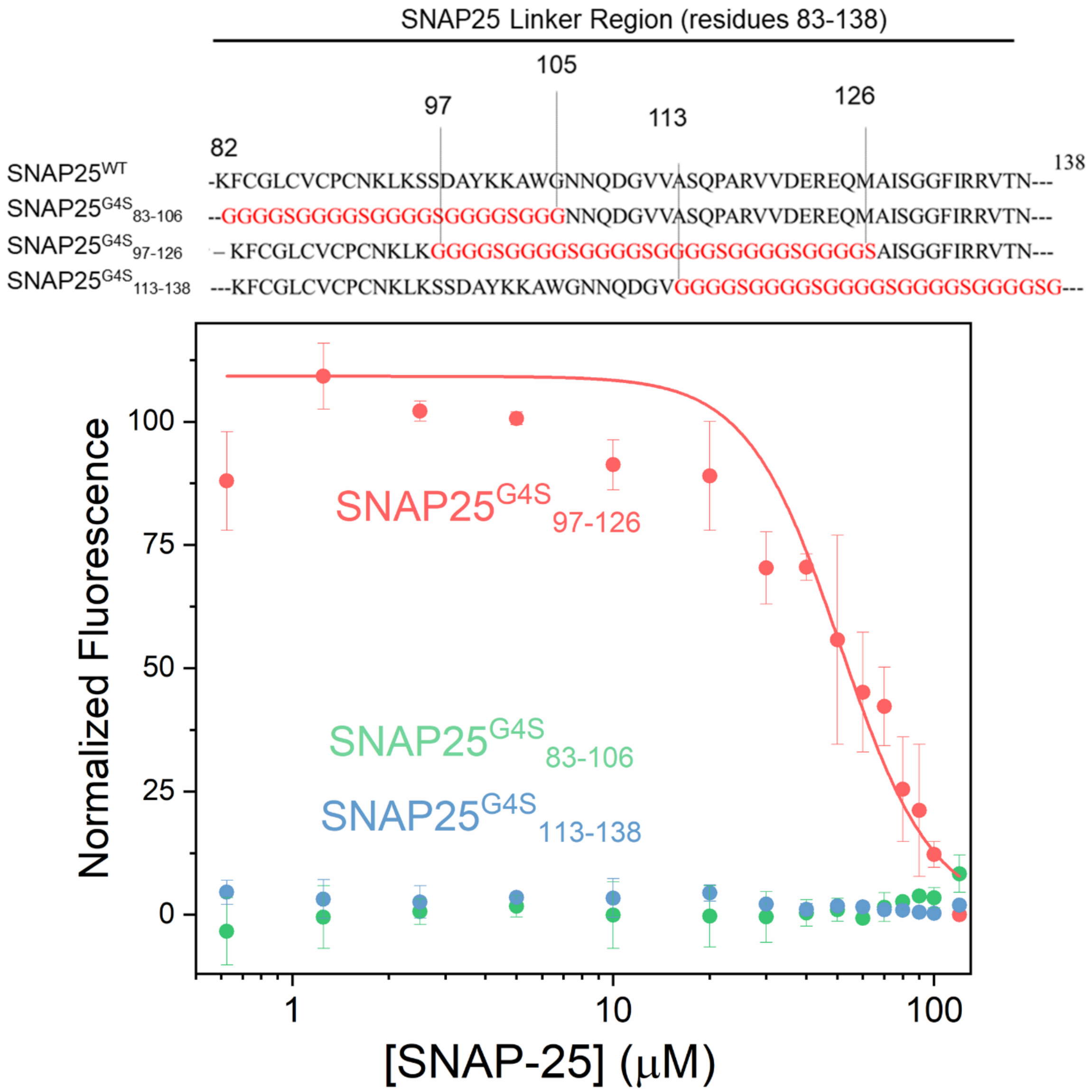


***Figure S8.*** Design of SNAP25 mutant that lacks the ability to bind Munc13. Based on our previous discovery (1) of direct binding between Munc13 and the SNAP25 linker region (residues 83-138), we generated and tested SNAP25 variants with portions of the linker region replaced by the GGGGS sequence ($SNAP25^{G4S}$) using Microscale Thermophoresis (MST) Analysis. Titration of full-length SNAP25 variants into AlexaFluor 660 labeled Munc13 a (using a C-terminal Halo tag) revealed that replacing either the N-terminal end (residues 83-106, green) or the C-terminal end (residues 113-138, blue) of the linker with GGGGS residues completely abolished the Munc13-SNAP25 interaction. However, replacing the middle portion (residues 97-126) had no effect on binding. Considering that the N-terminal portion is palmitoylated and associated with the membrane, we selected the C-terminal variant ($SNAP25^{G4S}_{113-138}$) for further detailed functional analysis. Average and deviations from three independent experiments are shown.

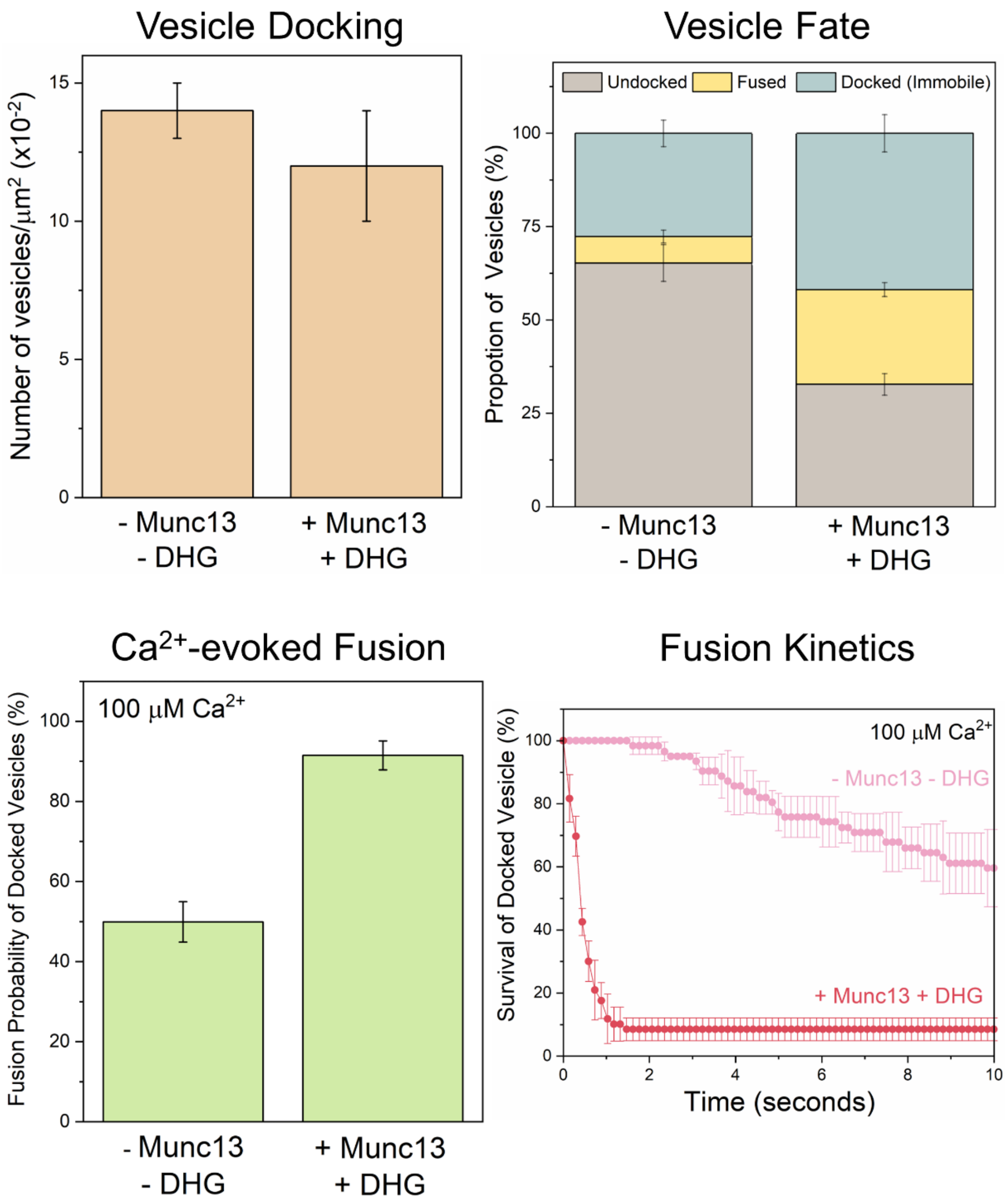


***Figure S9.*** We assessed the impact of the Munc18$^{DK}$ mutation, known to facilitate the formation of the Munc18-Syntaxin-VAMP2 'template' complex, on vesicle docking, priming, and $Ca^{2+}$-evoked fusion in the absence of Munc13/DHG. We observed similar total vesicle numbers and proportions converting to an immobile state, regardless of the presence or absence of Munc13/DHG. However, the probability and kinetics of $Ca^{2+}$-evoked fusion were markedly different between the two conditions when involving the Munc18$^{DK}$ mutant. The fusion probability was low in the absence of Munc13/DHG, and fusion kinetics was significantly slower. These findings highlight the collaborative role of Munc18 and Munc13 in generating stably docked vesicles that can be rapidly and reliably triggered for release by $Ca^{2+}$. The average values and standard deviations from three independent experiments (with ~150 vesicles per condition) are shown.

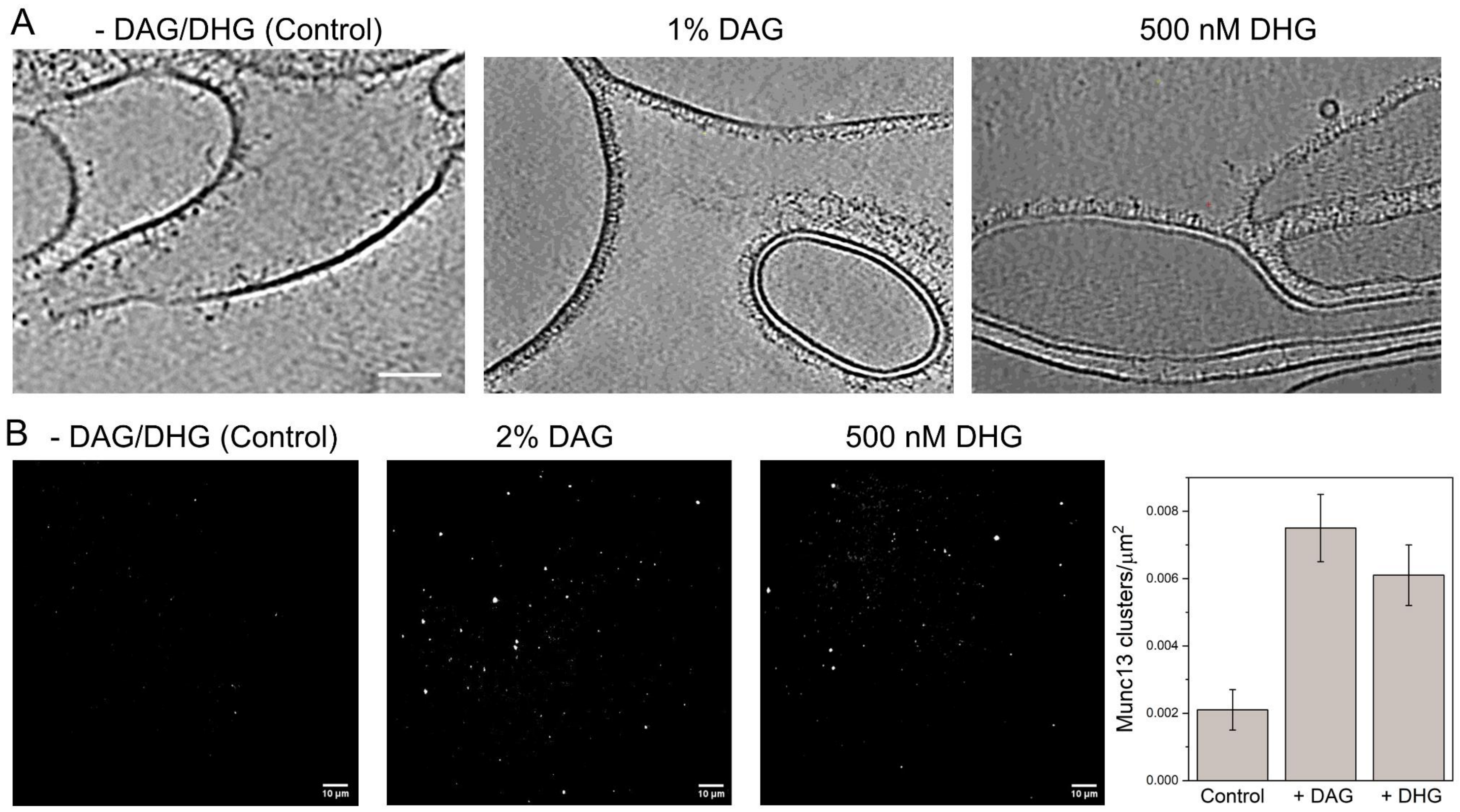


***Figure S10.*** Membrane association of Munc13 in the absence or presence of long-chain/short-chain DAG was assessed using cryo-electron microscopy (cryoEM) and TIRF microscopy. (A) For cryoEM experiments, Munc13 (250 nM) was incubated with pre-formed vesicles (DOPC/PIP2/DOPS 14/6/80) without or with DAG (2% in vesicles) or DHG (500 nM in solution) for 5 minutes in room temperature before freezing (2). cryoEM analysis reveals that the inclusion of 2% DAG increases the membrane association of Munc13 (middle panel) evidenced by brush-like electron density. We observed comparable Munc13 association and organization on lipid membranes in the presence of 500 nM DHG (right panel) in solution. (B) Representative TIRF images of Munc13 labelled with Alexa 488 on lipid bilayer membrane containing DOPC/DOPS/PIP2 (71/25/2) without or in the presence of 2% DAG in the membrane or 500 nM DHG added in solution. The inclusion of DAG or DHG increased the association and clustering of Munc13 on the membrane surface. Indeed, we observed comparable surface densities of the Munc13 clusters with DAG and DHG (3). Taken together, the data indicate that DHG serves as a suitable substitute for DAG.

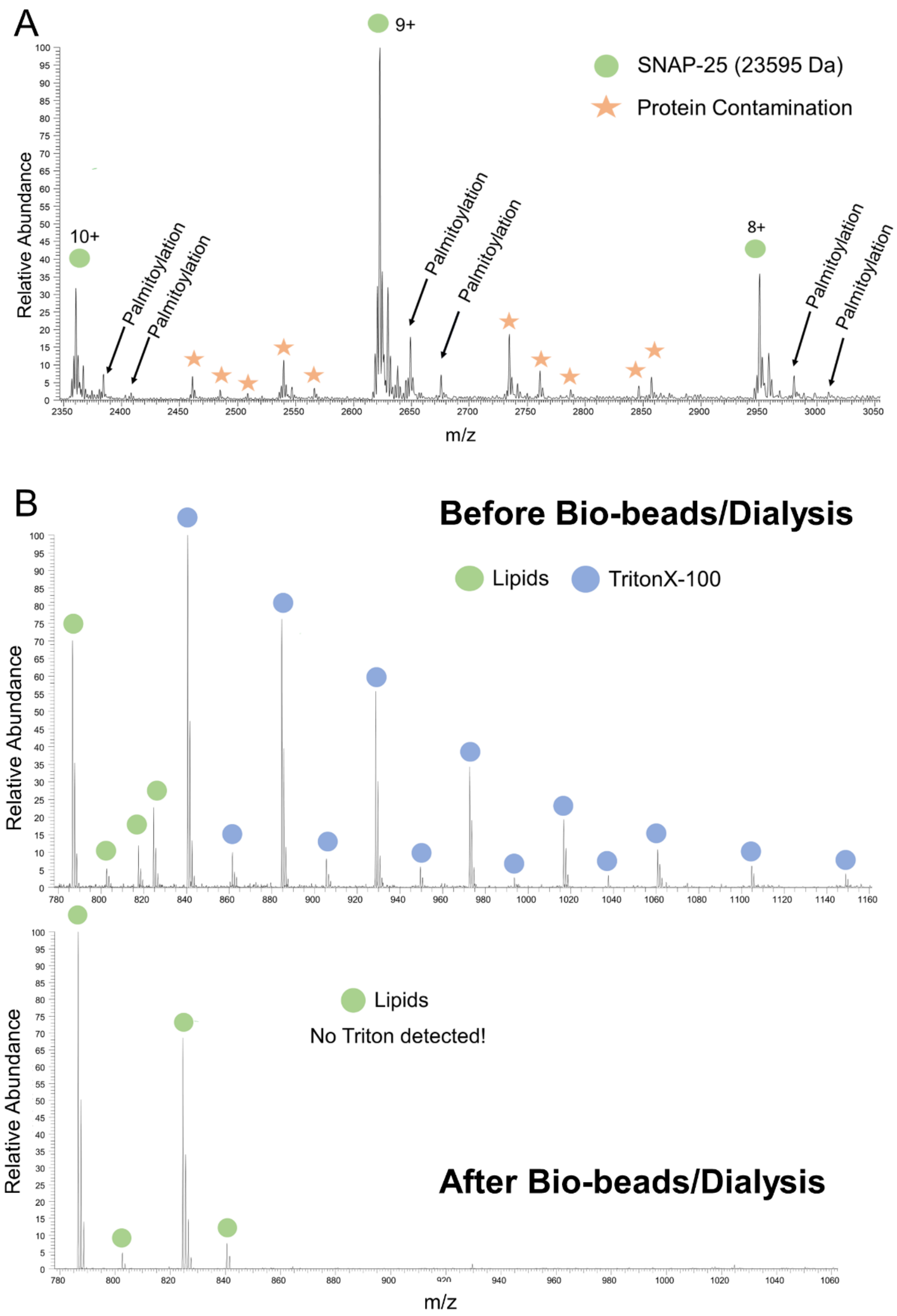


***Figure S11.*** Mass spectrometry analysis showing (A) palmitoylation of SNAP25 and (B) the effective removal of Triton X-100 following Bio-bead treatment and dialysis during Munc18/Syntaxin/SNAP25 liposome formation. The experimental details are described in Methods.

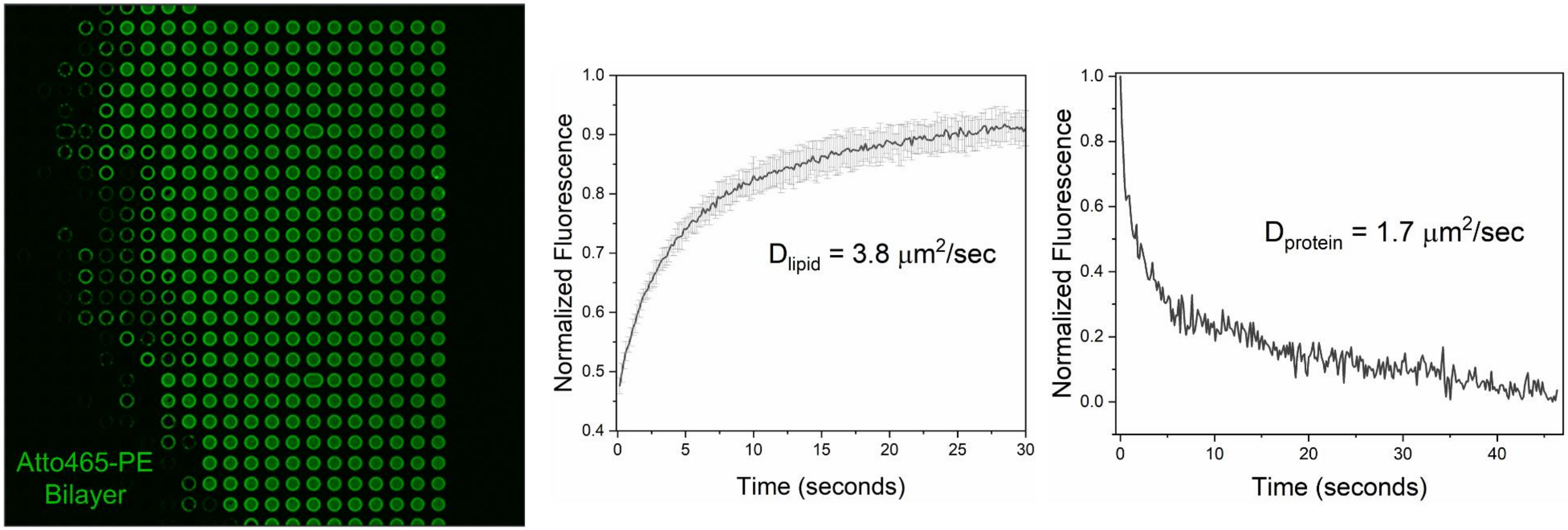


***Figure S12.*** The fluidity of the suspended lipid bilayer containing Munc18/Syntaxin and palymitoylated SNAP25 was confirmed using fluorescence recovery after bleaching (FRAP) analysis as described previously (4, 5). We used the fluorescence signal from ATTO465-PE (2%) included in the bilayer for lipid diffusion. We used Alexa568-NHS ester to label all proteins on the bilayer surface prior to FRAP analysis to estimate the protein diffusion on the suspended bilayer. FRAP measurements were analyzed on Wolfram Mathematica using a custom script to estimate the diffusion coefficient (4, 5).
